# *A Wolbachia pipientis* protein confers resistance to virus infection in *Drosophila melanogaster*

**DOI:** 10.64898/2026.09.21.753151

**Authors:** Thomas Leech, Bruno Taschke, Ralf Leslie Meilenbrock, Luke Stephen Tain, Pingze Zhang, Sebastian Grönke, Linda Partridge

**Author notes:** Corresponding authors: &.

## Abstract

The intracellular bacterium *Wolbachia pipientis* alters the biology of its arthropod hosts in many ways, and can increase resistance to RNA viruses in both *Drosophila* and mosquitoes. *Wolbachia*-induced pathogen blocking has generated much interest because of its potential to restrict insect vector transmission of human diseases caused by RNA viruses. However, the molecular mechanisms by which *Wolbachia* affects host viral resistance are still mostly elusive. We used *dilp2-3,5* mutant *Drosophila*, which are long-lived, but only in the presence of *Wolbachia*, to show that the presence of *Wolbachia* also increased the resistance of the mutant flies to infection with *Drosophila* C virus (DCV), relative both to mutants lacking *Wolbachia* and to wild type flies with and without *Wolbachia*. The insulin mutant flies had higher *Wolbachia* titres than wild type flies. By RNA-seq analysis of the *Wolbachia* transcriptome, we identified *Wolbachia* genes that were more strongly expressed in *dilp2-3,5* mutant flies. Ankyrin-domain-containing proteins were among the most strongly up-regulated and they are predicted to be secreted effector proteins. To address their effect on host physiology, we generated 4 transgenic fly lines each with inducible expression of a different genes encoding ankyrin-domain-containing proteins. Expression of 3 of these did not cause obvious effects, but expression of WD0754 at high levels severely shortened fly survival. Interestingly, however, chronic low-level induction of WD0754 increased the resistance of the flies to DCV infection. Proteomics analysis showed a robust, tissue-specific, anti-viral response upon WD0754 induction, and identified the NFkB-like IMD pathway as a potential mediator of the antiviral activity of WD0754. Consistently, WD0754 expression activated Relish, a key downstream transcriptional mediator of IMD signalling, while loss-of Relish blocked the antiviral effect of WD0754. In summary, we identified a *Wolbachia*-derived ankyrin domain containing protein that modulates host immunity through the IMD pathway.

**Authors’ summary:** *Wolbachia* is an intracellular bacterium that is able to proliferate in many insect species. Part of its evolutionary success is due to its ability to protect its host from infection with RNA viruses, but how *Wolbachia* does this is unknown. *Drosophila* that are mutant for insulin signaling are often long-lived, but only in the presence of *Wolbachia*. We found that flies lacking several of the *Drosophila* insulin ligands were protected against infection by *Drosophila* C virus (DCV), but again only in the presence of *Wolbachia*. A group of *Wolbachia* genes were upregulated in the *Drosophila* insulin mutants, suggesting that they could be responsible for the anti-viral protection. When we expressed one of these possible effectors, the ankyrin-domain-containing protein WD0754, in flies it could provide protection from DCV infection and induced upregulation of anti-viral immunity proteins. We further characterised this WD0754-induced anti-viral immunity and found that it required Relish, a well-known *Drosophila* immune transcription factor. This work marks an important step in understanding *Wolbachia*-induced pathogen blocking, especially in the context of *Wolbachia’s* use as a biological control agent for vector-borne, pathogenic RNA viruses of humans.

## Introduction

*Wolbachia pipientis* is an obligately intracellular bacterial symbiont, almost unrivalled in its ability to manipulate host physiology and the breadth of species it is able to infect [1]. The evolutionary success of this alpha-proteobacterium relies on the many and varied strategies it employs to modulate aspects of host biology, including the induction of reproductive parasitism [2] and increased resistance to RNA viruses [3], [4]). Understanding how *Wolbachia* protects its host from viral infection is important given its use as a control agent for insect-vector-borne viral diseases of humans [5], [6].

There are multiple ways in which *Wolbachia* could establish a host environment hostile to viruses [7]. *Wolbachia* infection reorganises membrane architecture [8] and the cytoskeleton [9],[10], [11], possibly interfering with viral entry into the cell. Especially well established is *Wolbachia’s* alteration of cholesterol and lipid metabolism and localisation [12] [13] [14]. These changes make the cell more suitable for *Wolbachia* infection, but simultaneously less hospitable for RNA viruses. *Wolbachia* infection also induces oxidative stress; strains that produce greater levels of reactive oxygen species (ROS) are better protected from infection [15]. Direct competition with *Wolbachia* for nutrients such as amino acids and iron may also make viral replication difficult. More recent work has suggested that the *Wolbachia* gene *RNase HI* contributes to virus blocking in the mosquito *Aedes aegypti* by degrading viral RNA [16]. In some cases, *Wolbachia* is also capable of protecting *D. melanogaster* from bacterial [17] and fungal [18] infections, suggesting induction of a broader immune response.

The evolutionarily conserved Insulin/Insulin-growth factor signalling (IIS) pathway is a key regulator of development, metabolism and longevity [19]. IIS also regulates innate immunity [20] and has been implicated in *Wolbachia*-dependent blocking of replication of dengue and Zika viruses in *Aedes aegypti*-derived cells [21]. *Wolbachia* also interacts with IIS in *Drosophila* [22]. Flies expressing a dominant-negative version of the *Drosophila* insulin receptor show a range of insulin mutant phenotypes that are reduced in the presence of *Wolbachia*, suggesting that the symbiont increases insulin signalling [22]. Consistent with this idea, mutant *dilp2–3,5* triple null flies, which completely lack 3 of the fly insulin-like peptides, exhibit strong mutant phenotypes including a substantial lifespan extension that is dependent on the presence of *Wolbachia* [23]. Thus, *Wolbachia* may rescue adverse effects of the strongly lowered IIS in these flies. *Wolbachia* did not affect insulin-signalling-related phenotypes, including lifespan, in the wild type control strain in either study. To discover if the interactions between *Wolbachia* and lowered insulin signalling extends to the virus-blocking effect of *Wolbachia*, we used the *dilp2–3,5* mutant to determine whether *Wolbachia* could also increase its resistance to viral infection, and to identify candidate *Wolbachia* proteins involved.

We show that *dilp2–3,5* mutant flies, which carry higher levels of *Wolbachia* than wild type flies, are resistant towards infection with *Drosophila* C virus (DCV). RNA-seq profiling of the *Wolbachia* transcriptome identified candidate *Wolbachia* genes with increased abundance in the mutant. Ankyrin-domain containing proteins, which are potential secreted *Wolbachia* effector proteins, were among the most strongly induced *Wolbachia* genes. We tested the function of 5 of them by expressing them directly in wild type flies. One ankyrin-domain containing *Wolbachia* protein, WD0754, drastically reduced lifespan when expressed at high levels. However, low-level induction of WD0754 improved survival upon DCV infection and reduced DCV titres. By performing a tissue-specific proteomic analysis, we identified the NFkB-like IMD pathway as a potential mediator of the antiviral function of WD0754. WD0754 expression caused cleavage and nuclear accumulation of Relish, indicating increased activity of the key downstream transcription factor of IMD signalling. Relish function was essential for the antiviral effects of WD0754. We have thus identified a *Wolbachia* protein that can enhance host immunity and protect against viral infection via activation of the IMD pathway, and that does so specifically in the context of reduced insulin signalling.

## Results

### *Wolbachia* increases viral resistance of *dilp2, 3-5* mutants

To test whether *Wolbachia* can protect *dilp2, 3-5* mutants from viral infection, we measured survival of female mutants with or without *Wolbachia* upon infection with *Drosophila* C virus (DCV). The outbred *wDah* wild type strain, which naturally carries *Wolbachia*, was used as the wild type control. *wDah* flies were treated with Tetracycline to generate the *Wolbachia*-free *wDahT* control strain. The mutant *dilp2-3,5* alleles were backcrossed into the *wDah* and *wDahT* genetic background generating *Wolbachia*-positive *wDah; dilp2-3,5* and *Wolbachia*-negative *wDahT; dilp2-3,5* flies. *Wolbachia* status of these flies was confirmed by PCR (**Fig S1A**).

*wDahT; dilp2-3,5* mutants without *Wolbachia* were slightly more susceptible than *wDahT* control flies to DCV challenge (**Fig 1A**), while *wDah; dilp2-3,5* mutants with *Wolbachia* showed higher survival upon DCV infection than mutants without *Wolbachia* and the corresponding wild type controls with and without *Wolbachia* (**Fig 1A**). A repeat of this experiment showed a similar pattern, but without a significant difference between *wDahT*; *dilp2-3,5* flies without *Wolbachia* and wild types (**Fig S1A**). *Wolbachia* thus increased the resistance of the insulin mutant flies to DCV infection.

**Figure 1:**
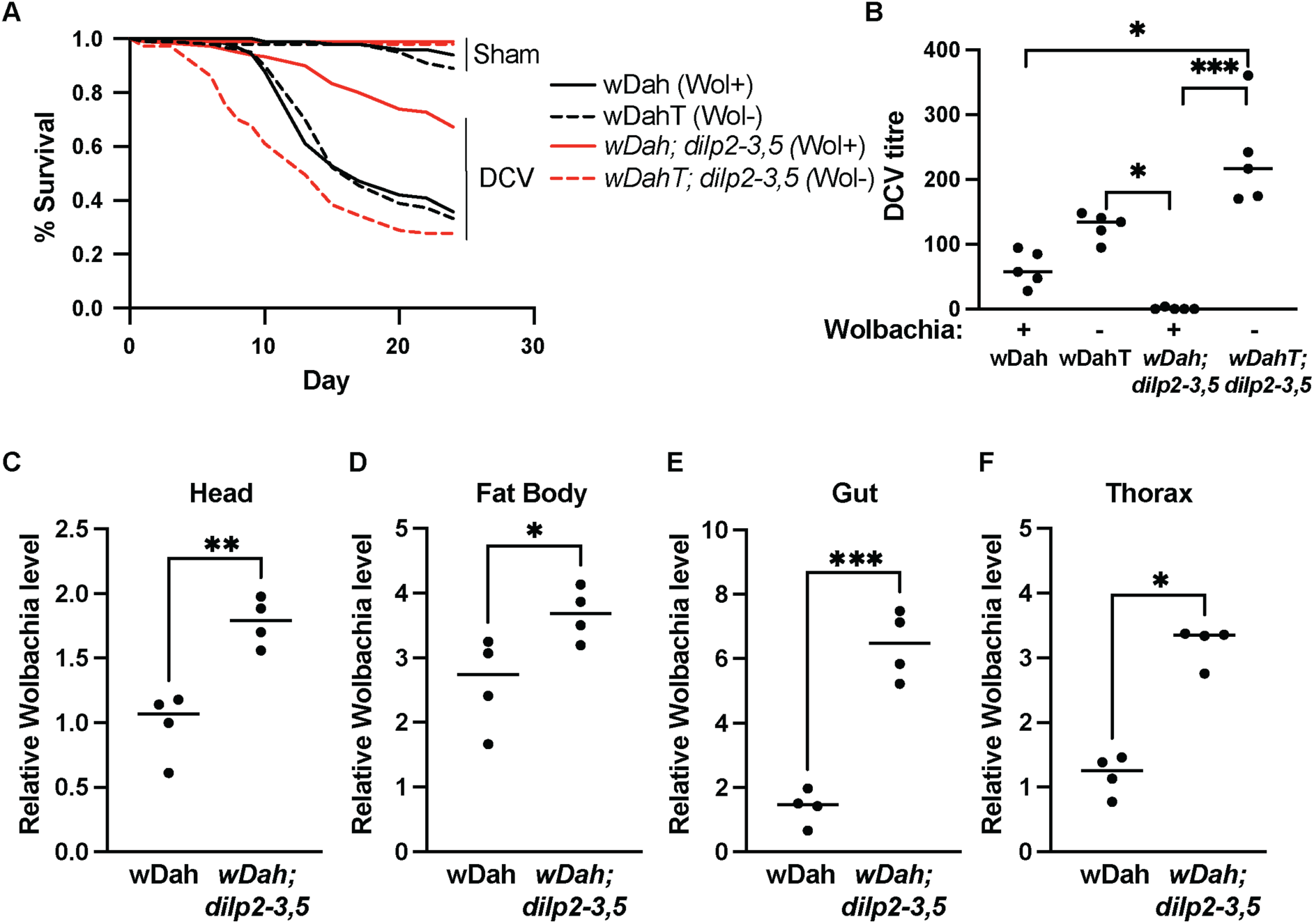
*Wolbachia* increases viral resistance of *dilp2-3,5* mutants. **(A)** Survival of female wild type wDah and *dilp2-3,5* mutant flies with (Wol+, full line) and without (Wol-, dotted line) *Wolbachia* upon infection with *Drosophila* C virus (DCV). *Wolbachia*-positive wDah; *dilp2-3,5* mutants were significantly more resistant towards DCV infection than *Wolbachia*-negative wDahT, *dilp2-3,5* mutants (p < 0.0001, Gehan-Breslow-Wilcoxon test at day 20, P-values were adjusted for multiple comparisons using Bonferroni correction). *Wolbachia*-negative wDahT; dilp2-3,5 mutants showed significantly reduced resistance towards DCV infection compared to wild type control flies (p < 0.005). **(B)** RT-qPCR analysis of DCV titre in *wDah* wild type and *wDah; dilp2-3,5* mutant females with and without *Wolbachia*. Kruskal-Wallis test using Dunn’s test for comparisons. n=5 groups of 20 pooled females. Titre was measured 72 hours post infection. (**C-F**) *Wolbachia* titre measured by RT-qPCR in the (**C**) head, (**D**) fat body, (**E**) gut and (**F**) thorax of 10-day-old wild type wDah and *dilp2-3,5* mutant flies. A probe targeting the *Wolbachia* surface protein (*wsp)* gene was used and normalized using *Rpl32*. n=4 groups of 20 flies. Unpaired t test. *p < 0.05, **p < 0.01, ***p < 0.005.

We next measured DCV titres by qRT-PCR post infection. Consistent with the survival data, *wDah; dilp2-3,5* flies with *Wolbachia* had extremely low DCV levels, whilst *wDahT; dilp2-3,5* mutants without *Wolbachia* showed the highest DCV levels (**Fig 1B**), suggesting they were especially sensitive to DCV infection. Surprisingly, DCV resistance was not significantly different between *wDah* and *wDahT* control flies. There was a non-significant trend for lower DCV titres in *wDah* compared to *wDahT* flies, suggesting that *Wolbachia* may also have a virus blocking effect in control flies, which was not strong enough to affect survival upon DCV infection (**Fig 1B**). As anti-viral resistance is associated with high levels of the symbiont [24][25] we determined if *Wolbachia* was present at higher densities in insulin mutant flies, and found that they indeed had significantly higher levels of *Wolbachia* than wild type flies in all tested tissues (**Fig 1C-F**). Together these results indicate that *dilp2-3,5* mutants, which carry higher levels of *Wolbachia* than wild type flies, are protected from DCV infection, while *dilp2-3,5* mutants without *Wolbachia* and wild type flies are more sensitive.

### *Wolbachia* Ankyrin domain containing proteins are upregulated in *dilp 2, 3-5* mutants

In order to understand how *Wolbachia* confers virus resistance in *wDah; dilp 2-3,5* mutants, we profiled the *Wolbachia* transcriptome using RNA-seq in *wDah* and *wDah; dilp 2-3,5* mutant flies with *Wolbachia*. This dataset was first published with an emphasis on changes in the host, rather than *Wolbachia* [26]. When normalised to the total *Wolbachia* genome, we detected 729 *Wolbachia* transcripts, of which 102 were significantly differentially regulated between the Wolbachia transcriptomes of *dilp2-3,5* mutants and wild type flies. Of these, 71 were regulated in the thorax, 28 in the gut, 42 in the fat body, and none in the head (**Fig 2A, SI 1**).

**Figure 2:**
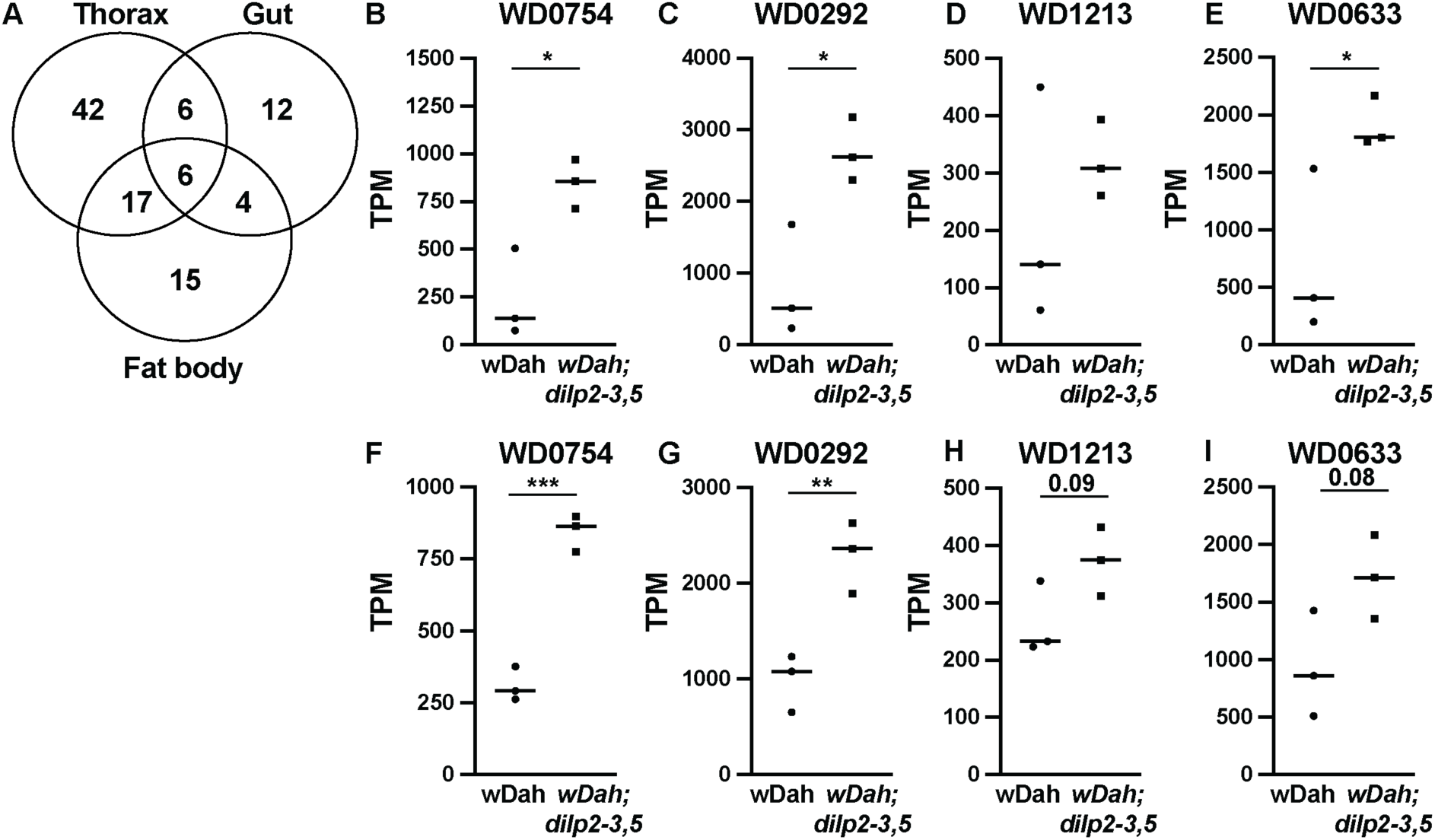
Identification of ankyrin-domain containing *Wolbachia* proteins with increased abundance in virus-resistant *dilp2-3,5* mutant flies. *Wolbachia* genes with differential abundance between *wDah; dilp2-3,5* and *wDah* control females were identified by RNAseq [26] (**A**) Venn diagram outlining the number of Wolbachia genes with significant differential abundances in the thorax, gut and fat body of *wDah; dilp2-3,5* mutants compared to *wDah* flies. (B-I) Expression of ankyrin-domain containing *Wolbachia* proteins in (B-E) the thorax and (F-I) the fat body of wt and *dilp2-3,5* mutant flies. Expression levels of WD0754 (B, F), WD0293 (C, G) and WD0633 (E, I) were significantly higher in *dilp2-3,5* mutants compared to control flies, while a similar but non-significant trend was observed for WD1213 (D, H). Expression is based on the RNA seq data and given as transcripts per million (TPM). Mean with SEM. *p < 0.05, **p < 0.01, ***p < 0.005, unpaired t test.

GO analysis did not reveal enrichment for specific functional pathways in any tissues, probably in part because around 35% of the differentially regulated *Wolbachia* genes are still uncharacterised. In the thorax (**Fig 2B-E**) and fat body (**2F-I**), however, we observed that 2 of the most significantly up-regulated Wolbachia transcripts in *dilp2-3,5* mutant flies encode the ankyrin domain containing proteins WD0754 (**Fig 2B, F)** and WD0292 (**Fig 2C, G**). The transcripts of two further ankyrin-domain containing proteins WD1213 (**Fig 2D, H)** and WD0633 (**Fig 2E, I)** also showed a trend for upregulation in the thorax and fat body. Ankyrin domains are often involved in protein-protein interactions [27] and, whilst they are relatively rare in bacterial genomes, they are occasionally present in intracellular bacteria, where they can interfere with host biology [28]. Ankyrin domain containing proteins can be secreted and, interestingly, WD0754, WD0292 and WD0633 were predicted by the EffectiveELD algorithm to be secreted proteins based on the presence of eukaryotic-like domains (ELD)[29]. ELDs are also found in eukaryotic genomes, and bacterial proteins containing ELDs may act as effectors in the host cell [29].

### Expression of the ankyrin domain containing protein WD0754 causes tissue-specific toxicity in *Drosophila*

To investigate possible roles of the *Wolbachia*-derived ankyrin-domain containing proteins in host biology, particularly health and immunity, we generated *Drosophila* lines that expressed the ankyrin-domain containing proteins WD0754, WD0292, WD1213 and WD0633 in an inducible manner. The constructs were codon optimized for *Drosophila* and contained a *Drosophila* Kozak sequence, to allow robust expression in the fly. Efficient induction of transgene expression was verified for all four transgenic lines by Q-RT-PCR (**Fig S2A-D**). To address whether the ankyrin domain containing *Wolbachia* proteins affect fly physiology, we first measured lifespan of female flies upon adult-onset ubiquitous expression of the transgenes using the *daughterless* Gene Switch (*da*-GS) driver line both in the presence (**Fig S3A-D**) and absence (**Fig 3A-D**) of *Wolbachia*.

**Figure 3:**
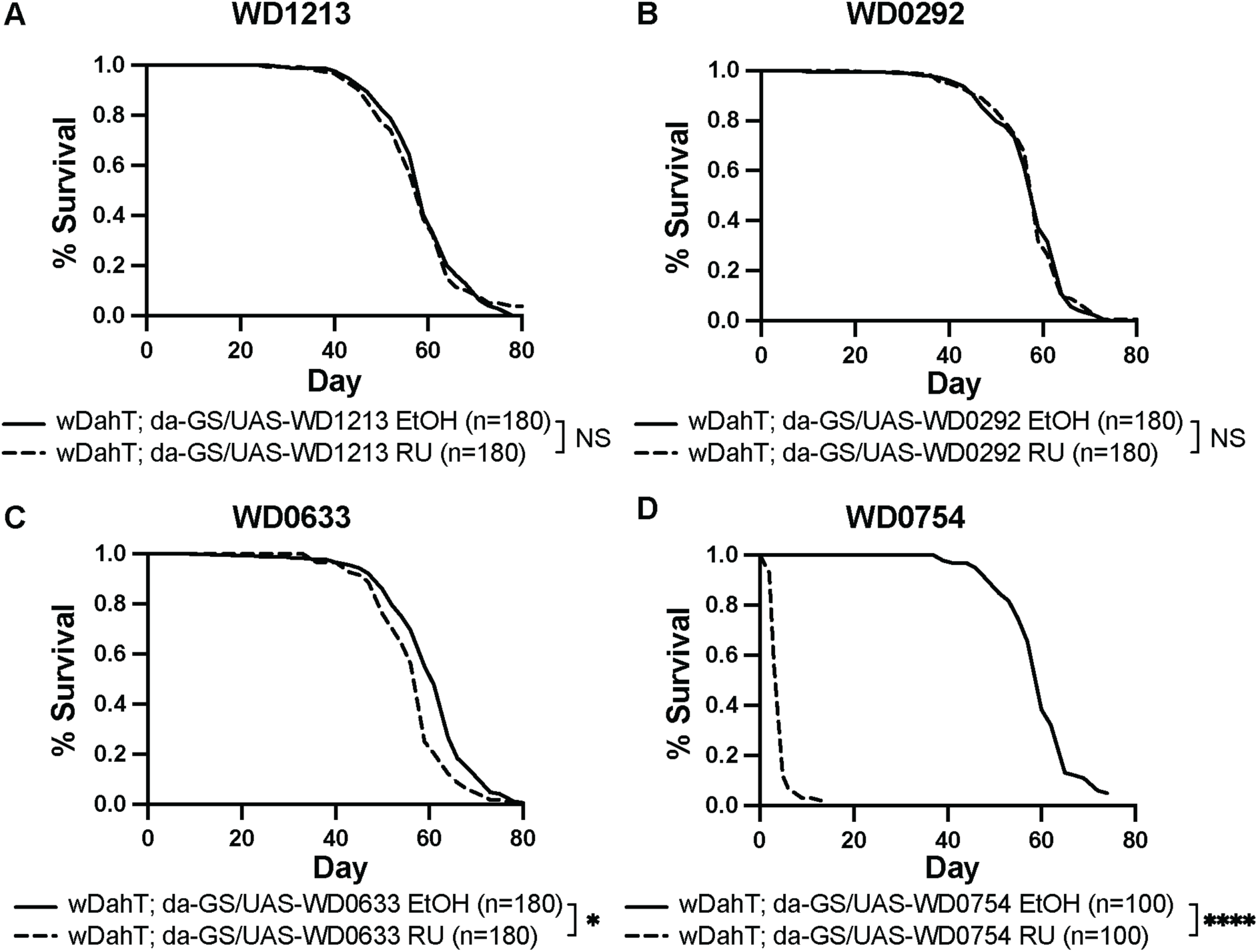
Overexpression of WD0754 drastically shortens lifespan. **(A-D)** Survival of female flies ubiquitously overexpressing the ankyrin-domain containing *Wolbachia* proteins (**A**) WD1213, (**B**) WD0292, (**C**) WD0633, (**D**) and WD0754 under the control of the da-GS driver. Expression of the transgene was induced with 200µM RU486 (RU). Non-induced control animals only received the carrier EtOH (EtOH). While overexpression of WD1213, WD0292 and WD0633 had only minor or no effects on fly survival, overexpression of WD0754 dramatically shortened lifespan (*: p < 0.05, ****: p < 0.0001, log-rank test). Number in brackets indicate the number of females used per treatment. All flies were free of *Wolbachia*. The corresponding lifespan data for Wolbachia-positive flies are shown in Fig S3.

Expression of WD1213 (**Fig 3A, S3A),** WD0292 (**Fig 3B, S3B)** and WD0633 (**Fig 3C, S3C)** had only minor effects on survival. In contrast, ubiquitous adult-onset expression of WD0754 caused a drastic decrease in lifespan from a median of 59 days in EtOH fed control flies, to 3 days upon induction with 200µM RU (**Fig 3D, S3D**). The lifespan shortening effect of WD0754 expression was independent of the presence of *Wolbachia,* suggesting that WD0754 directly acts on the host and not indirectly via affecting Wolbachia physiology. WD0754-expressing flies also showed strongly reduced egg-laying (**Fig S4A**), decreased food intake (**Fig S4B**) and reduced body weight (**Fig S4C**). Thus, ubiquitous expression of WD0754 was extremely toxic for adult flies. Intriguingly, toxicity of expression was tissue-specific. Expression of WD0754 specifically in the enterocytes of the gut or in the flight muscle, using 5966-GS and act88F-GS drivers respectively, caused only a very small (**Fig 4A**) or non-significant (**Fig 4B**) decrease in lifespan. In contrast, neuron-specific expression of WD0754 using the elav-GS driver shortened lifespan to the same extent as ubiquitous expression, with a median lifespan of around 3 days (**Fig 4C**). We also tried to express WD0754 in the adult fat body using the Larval serum protein 2 Gene Switch driver (Lsp2-GS), but larvae on control food did not develop into adult flies. Leaky expression alone of the Lsp2-GS driver during development in combination with the WD0754 transgene was sufficient to kill the flies, suggesting that WD0754 is also very toxic when expressed in the fat body. In summary, the *Wolbachia*-derived ankyrin domain protein WD0754 is highly toxic *in vivo* in a tissue dependent manner.

**Figure 4:**
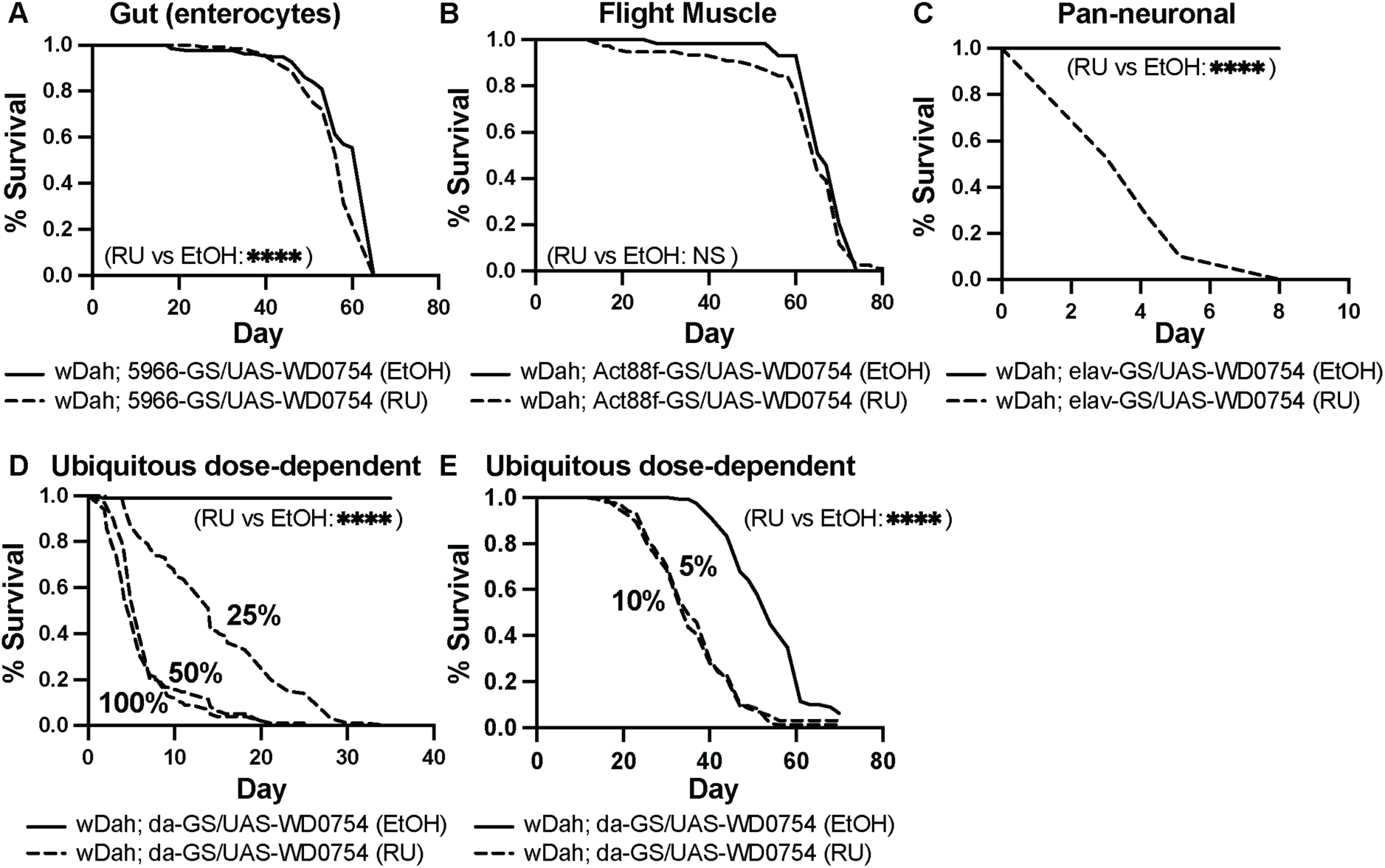
Overexpression of WD0754 causes toxicity in a tissue-specific and dose-dependent manner. **(A-B)** Overexpression *of* WD0754 *in* (***A***) *gut* enterocytes and (***B***) flight muscles using 5966-GS and Act 88f GS, respectively, did not strongly affect the survival of female flies. In contrast, overexpression of WD0754 using the pan-neuronal elav-GS driver line (***C***) dramatically reduced survival of female flies. Expression was induced using 200µM RU486. (***D-E***) Survival of female flies ubiquitously overexpressing WD0754 under the control of the da-GS driver with reduced concentrations of the inducing agent RU486 (100% = 200µM, 50% = 100µM, 25% = 50µM in (***D***) and 10% = 20µM, 5% = 10µM RU486 in (***E***)). Toxicity of WD0754 expression was dose-dependent, and flies that received 25% of the inducing agent were longer-lived than flies receiving 50% or 100%. Reducing RU486 concentration to 10% and 5% further increased lifespan compared to the higher doses, while there was no difference between the two lower doses. All flies were *Wolbachia* positive females. ****: p < 0.00001, log-rank test. n = 140 flies for (***A***), n = 80 for induced and n = 60 for control in (***B***), n = 100 for induced and n = 80 for control in (***C***), n = 100 for (***D***), n = 160 for (***E***).

The normal degree of exposure of fly cells containing *Wolbachia* to WD0754 and other *Wolbachia* proteins is unknown. The strong toxicity that we found with direct expression may have been caused by unphysiologically high cellular levels of the WD0754 protein, and it prevented further analysis of a potential function of WD0754 in viral resistance. Therefore, we next addressed whether toxicity could be reduced by titrating down the level of the inducing agent RU486 in females expressing WD0754 under the control of the da-GS driver (**Fig 4D, E**). While a 50% reduction in RU levels did not change survival compared to the full dose, flies survived significantly longer on 25% RU, with median lifespan increasing from 3-4 days on the full dose to 14 days (**Fig 4D**). Titrating RU levels down to 10% further increased survival, with a median lifespan of 35 days, while a reduction to 5% RU levels had no additional effect (**Fig 4E**). Thus, toxicity of WD0754 was dose-dependent and animals survived for up to 35 days on the lower doses, allowing further physiological phenotyping.

### WD0754 protects flies from *Drosophila* C Virus infection

We next tested whether adult-specific ubiquitous overexpression of the ankyrin-domain containing *Wolbachia* proteins under the control of the da-GS driver could protect flies against infection with *Drosophila* C virus (DCV). Overexpression of WD0292, WD1213 and WD0633 did not significantly affect survival upon DCV infection (**Fig 5A-C**). WD0754 was also not efficient in protecting flies against DCV when transgene induction was initiated 72h before infection and these flies even died slightly earlier than non-induced control flies (**Fig S5A**).

**Figure 5:**
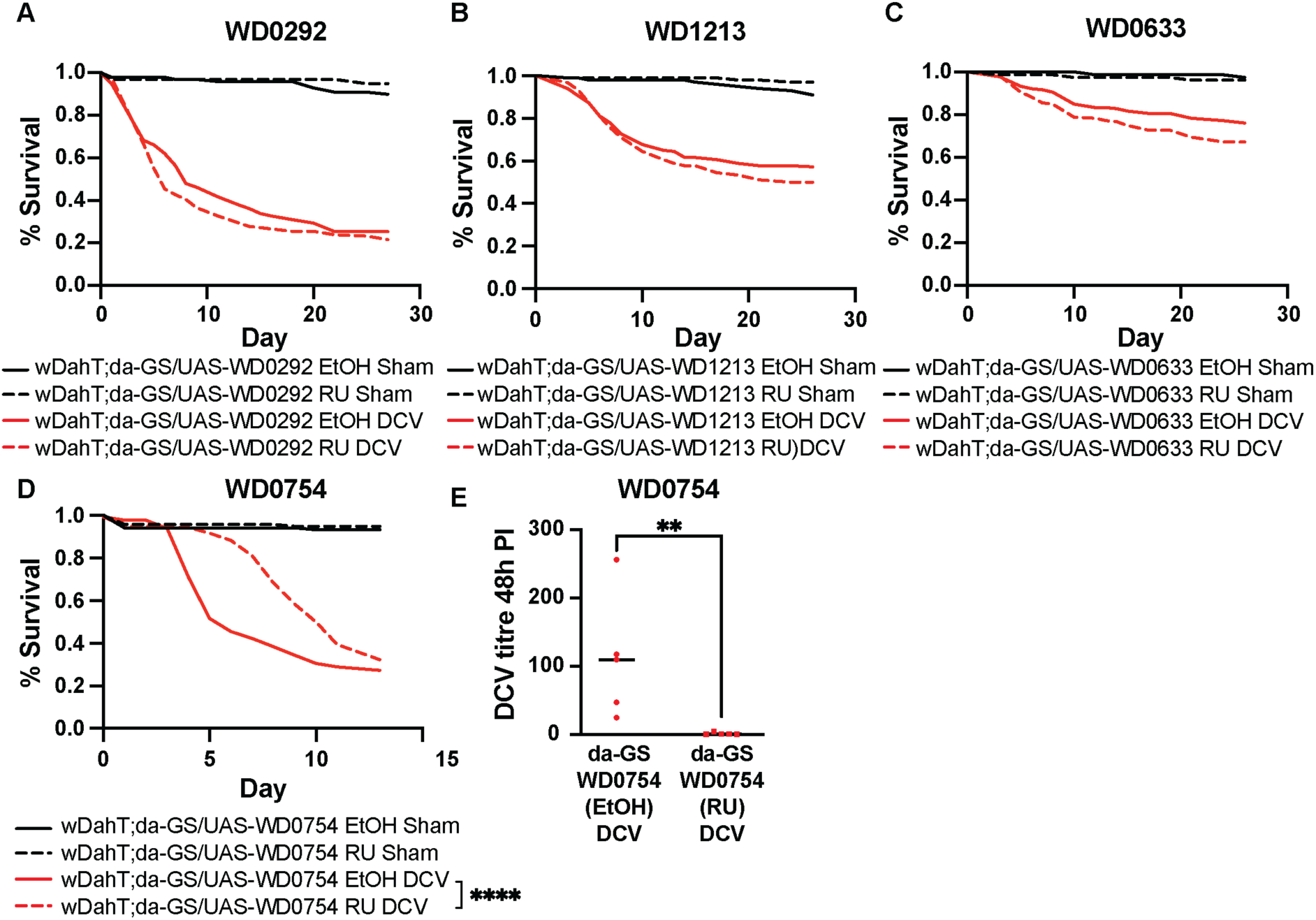
Expression of WD0754 protects flies from *Drosophila* C virus infection. **(A-C)** Survival of female flies infected with *Drosophila* C virus was not significantly changed upon ubiquitous expression of (**A**) WD0292, (**B**) WD1213 and (**C**) WD0633. Flies were infected with DCV (red) or sham treated with sterile PBS (black). Induction of transgene expression was initiated 72h prior to infection by feeding flies with 200µM RU486 (RU). Non-induced control flies received only the carrier EtOH (EtOH). (**D**) Induction of WD0754 transgene expression 12h before DCV infection with 10 µM RU486 significantly improved the survival of female flies (p < 0.00005, Gehan-Breslow-Wilcoxon. n=120 for sham and 180 for DCV infected flies in (**A-D**). (**E**) DCV titre as measured by RT-qPCR 48h post infection was significantly reduced upon WD0754 overexpression (**: p < 0.005, Mann-Whitney test. n = 6 groups of 20 pooled females. da-GS was used to drive transgene activation ubiquitously in **A-E**. See Sup Fig. 5B & 5C for an independent verification of **D & E**, respectively.

However, there was a strong interaction between survival of the injury during infection and expression of the WD0754 transgene (**Fig S6A**). This is evident from the induced sham controls, which died shortly after pricking, presumably due to injury (**Fig S5A**). This finding suggests that prolonged expression of WD0754 before infection interferes with the flies’ response to injury. To overcome this effect, WD0754 transgene induction was initiated only shortly (12h) before infection and was maintained post-infection. In contrast to the prolonged expression protocol pre-infection, shortening the time of WD0754 expression before infection to 12h had no detrimental effect on recovery from injury (see the induced sham controls in **Fig 5D**, **S5B**). However, induction of WD0754 increased the resistance of flies towards DCV infection, with a ∼2.5x fold increase in median survival post-infection (**Fig 5D**, **S5B**). This was also mirrored in DCV titres, which were significantly lower in WD0754-expressing flies 48h post-infection (**Fig 5E, S5C**). In summary, expression of WD0754 under conditions where it did not interfere with wound healing protected flies against DCV infection by repression of viral replication.

### WD0754 expression induces a proteome characterised by upregulation of anti-viral proteins

To address how WD0754 expression affects viral resistance, we measured changes in the fly proteome by mass spectrometry in the head, fat body and gut upon ubiquitous expression of WD0754 for 24h and 72h (**Fig 6 and SI 2**). In total 6122 proteins were detected across all 3 tissues, of which 361 and 2983 were significantly regulated at 24h (**Fig 6A**) and 72h (**Fig 6B**), respectively. We first focused on the acute response of the proteome towards WD0754 expression, as it might indicate which processes are directly affected by WD0754. While there were no significantly regulated proteins detected in the fat body after 24h of WD0754 induction, 31 and 336 proteins were significantly regulated in the head and gut, respectively **(Fig 6A)**. Gene ontology analysis identified GO terms associated with defence response to virus and innate immunity to be enriched in the head upon short-term induction of WD0754 (**Fig 6C**). Consistently, the anti-viral immune response proteins Vago and Vir-1 and the antimicrobial peptides Attacin-B (AttB) and Attacin-A (AttA) were among the strongest upregulated proteins in the head after 24h (**Fig 6D** and **SI 2**) and this upregulation persisted 72h after transgene induction (**Fig 6E**). The protein encoded by the CG32368 gene, which has previously been shown to be upregulated upon infection with a variety of viruses [30], [31], [32], [33], was also upregulated in the head after 24h and 72h of WD0754 induction (**Fig 6 D, E**). Antiviral proteins were also upregulated in the gut after induction of WD0754 expression (**Fig 6F, G**). This includes AGO2 and Vago and AGO2, Vago and vir1, which were induced in the gut after 24h (Fig 6F) and 72h (Fig 6G) of WD0754 expression, respectively. While no proteins were significantly regulated in the fat body after 24h of WD0754 induction, there was a non-significant trend for Vago to be upregulated (**Fig 6H**), which became significant after 72h (**Fig 6I**). Another important immune regulator induced in the fat body by WD0754 after 72h, was Relish (Rel), a NF-kB-like transcription factor and key transcriptional effector of the IMD pathway (**Fig 6I** and **SI 2**) [34]. Other immune-related proteins significantly regulated at this timepoint included Dif, the transcription factor responsible for carrying out the Toll-specific transcriptional programme [35] and PGRP-SB1, which is a catalytic peptidoglycan recognition protein. Notably, PGRP-SB1 is abundantly secreted into the haemolymph upon Imd pathway activation in the fat body [36]. Diedel, a negative regulator of the IMD pathway that is upregulated in response to some virus infections [30], was also strongly upregulated at this timepoint. While some proteins like Vago and vir-1 were upregulated in several tissues, the response to WD0754 induction was also tissue-specific (**Fig 6A, B**), as can be seen by the different GO terms enriched in the head (**Fig S6A**), gut (**Fig S6B**) and fat body (**Fig S6C**) after 72h. Notably, in the head there was strong down-regulation of proteins involved in translation 72h after WD0754 induction (**Fig S6A** and **SI 2**), which is one proposed mechanism responsible for pathogen blocking [37].

**Figure 6:**
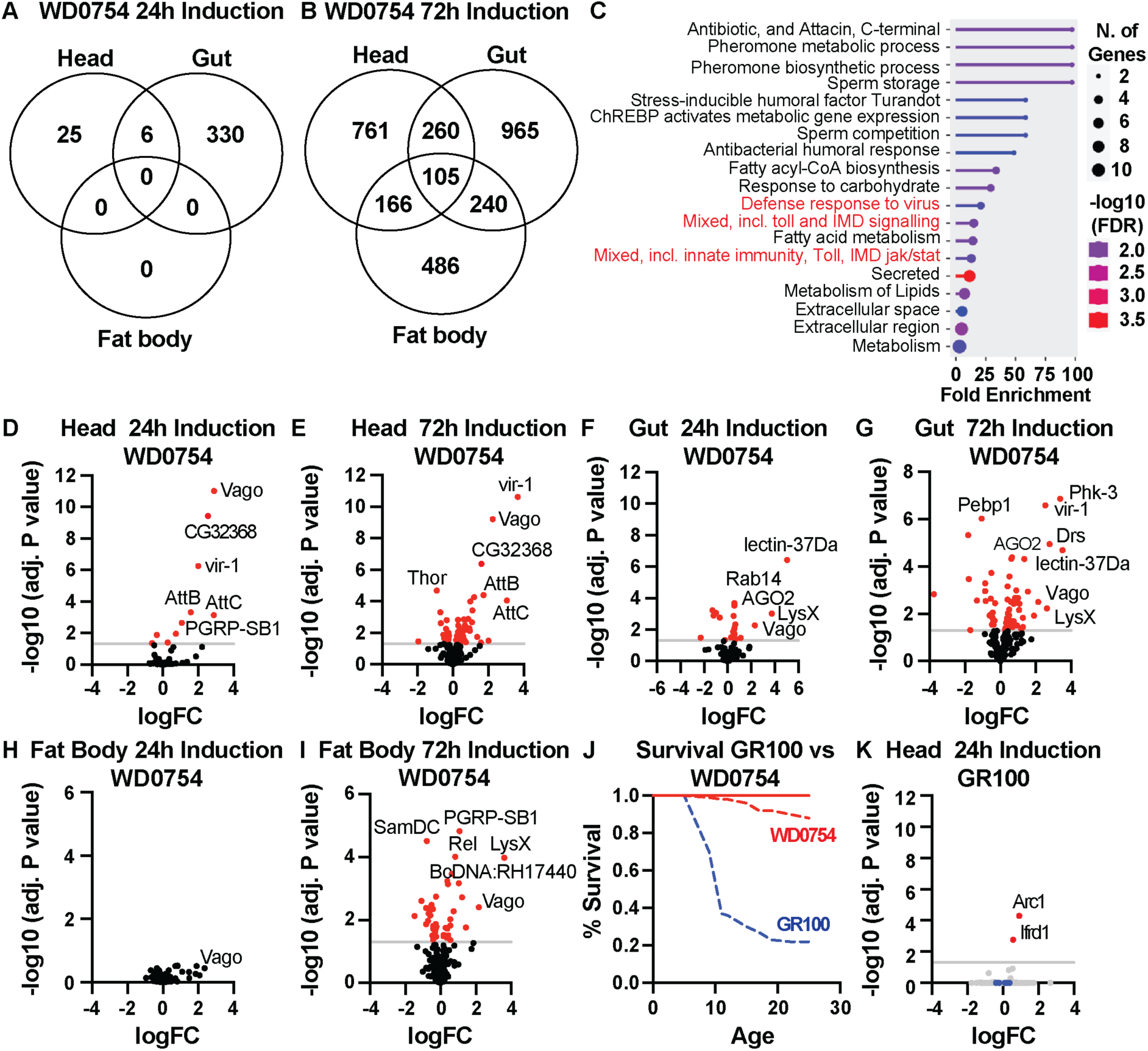
WD0754 expression induces upregulation of a tissue-specific anti-viral protein response. Changes in the head, gut and fat body proteome in response to ubiquitous WD0754 overexpression were measured via mass spectrometry. (**A**-**B**) Venn diagrams depicting significantly regulated proteins in head, gut and fat body after (**A**) 24 hours and (**B**) 72 hours of WD054 transgene induction using the ubiquitous da-GS driver line and 50µM RU486. (wDah; da-GS/UAS-WD0754 RU vs wDah; da-GS/UAS-WD0754 EtOH) (**C**) Gene ontology enrichment analysis of proteins differentially regulated in the head 24h post-transgene induction. GO terms associated with immunity are highlighted in red. (**D-I**) Volcano plots of proteins related to immunity in (**D-E**) head, (**F-G**) gut and (**H-I**) fat body 24h (**D, F, H**) and 72h (**E, G, I**) post WD0754 transgene induction. (**J**) Survival of female flies expressing WD0754 or GR100 under control of the ubiquitous da-GS driver. (**K**) Volcano plot of all detected proteins upon 24h induction of the toxic GR100 protein in fly heads using the ubiquitous da-GS driver line and 50µM RU486 (wDah; da-GS/UAS-GR100 RU vs wDah; da-GS/UAS-GR100 EtOH). In contrast to WD0754, expression of GR100 did not induce immune-related proteins. Blue dots indicate from left to right the immune-related proteins regulated by WD0754 expression: CG32368, Vir1, Vago, PGRP-SB1, AttA and AttC levels. Red dots in (**D-J**) indicate significantly regulated proteins (adjusted P < 0.05, n=6 biological replicates for **D-I)** and n=5 biological replicates for **J**. Every replicate contained 10 pooled tissues).

The immune induction in response to expression of WD0754 could have been the product of expressing any heterologous *Wolbachia* gene in the fly. We therefore generated proteomic profiles of the heads of flies expressing WD0292, WD0633 and WD1213 for 72h. In contrast to WD0754, induction of WD0292 and WD0633 did not cause any significant changes in the fly head proteome, whereas only 5 proteins were significantly upregulated upon expression of WD1213 (**Sup Fig 7**). These 5 proteins were different from the proteins regulated by WD0754 and were not associated with antiviral activity, demonstrating that the induction of an anti-viral protein response is specific to WD0754 and that this function is not shared by the other ankyrin-domain-containing protein we tested.

At high expression levels WD0754 is very toxic and the observed immune response might just be an unspecific response of the fly’s immune system to damage. In that case, other toxic proteins should induce a similar immune response in the proteome. To address this, we measured proteomic changes in the head upon ubiquitous expression of the toxic dipeptide repeat protein GR100 [38], which based on fly survival (**Fig 6J**) was even more toxic than WD0754. Despite the higher toxicity, GR100 expression did not cause an immune response in the proteome (**Fig 6K**), indicating that the immune induction is specific to WD0754 and not a general response towards toxic proteins.

### Relish is activated by WD0754 and essential for its antiviral protection

Vago, which is induced by WD0754 expression across tissues, is under the control of the NF-kB-like transcription factor Relish, at least in *Culex* [39]. In addition, other regulated proteins like Attacin B and C and PGRP-SB1 are also under transcriptional control of Relish in *Drosophila* [40], [36] and Relish itself was upregulated in the fat body upon long-term induction of WD0754 (**Fig 6I**). Thus, WD0754 might increase viral resistance by activating Relish signalling. Relish is the key downstream transcription factor of the IMD pathway, the prime immune response pathway in *Drosophila*. Upon pathway activation, Relish is cleaved in the cytosol and the activated fragment translocates to the nucleus to induce the transcription of target genes. In order to address whether WD0754 activates Relish, we first determined the subcellular localization of Relish by immunohistochemistry in young fat body cells using an αRelish antibody (**Fig 7A**). Relish was mostly located in the cytoplasm in control flies and under non-induced conditions. In contrast, expression of WD0754 for 24h caused a significant accumulation of Relish in the nucleus (**Fig 7B**), indicative of Relish activation. To further confirm this, we conducted Western blots on whole flies after 24h of WD0754 induction using an αRelish antibody that detects both the full length and the cleaved variants of Relish (**Fig 7C**). While we did not detect a smaller Relish fragment in control and un-induced flies (**Fig 7C**), induction of WD0754 caused a significant accumulation of the smaller form of Relish, consistent with its increased translocation to the nucleus (**Fig 7D**). In summary, WD0754 expression induced cleavage and nuclear translocation of Relish, suggesting that WD0754 activates Relish.

**Figure 7:**
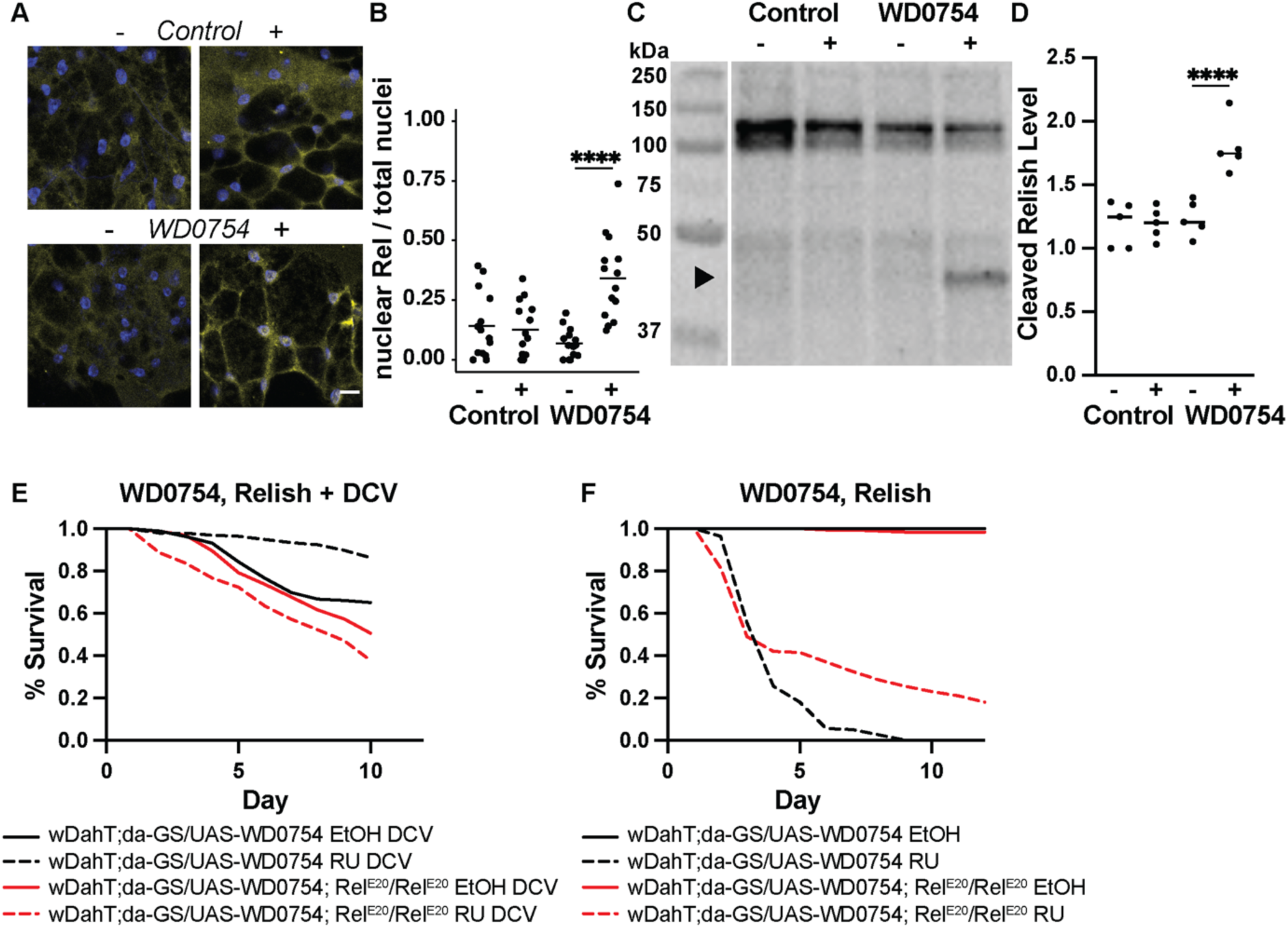
WD0754 induces protection against DCV via activation of Relish, a key downstream effector of IMD signalling. (**A-D**) Expression of WD0754 in the fat body results in nuclear accumulation (**A-B**) and activated cleavage (**C-D**) of the Relish transcription factor, a key downstream effector of IMD signalling. (**A**) Representative images of Relish protein localisation in fat body tissue of 7-day old female flies 24 post-induction of WD0754 expression. Control (wDahT; *da*-GS/+) and WD0754 (wDahT; *da*-GS/UAS-WD0754) flies were induced with 200µM RU486 (+) and as control with the carrier EtOH (-). Nuclei were stained with DAPI (blue) and relish staining is indicated in yellow. (**B**) Quantification of (**A**) Relish nuclear localisation, shows that induction of WD0754 causes a significant nuclear accumulation of Relish (****p<0.0005, linear mixed model, n = 16 tissue per treatment). (**C**) Representative western blot image and (**D**) the corresponding quantification of whole fly extracts probed with an αRelish antibody. Arrowhead indicates the size of the cleaved Relish49 fragment, which is significantly enriched 24h post-induction of WD0754 (n=5 replicates with 20 flies each, ****p<0.0005, One way-ANOVA). (**E**) The protective effect of WD0754 against DCV infection is dependent on relish function. Survival of female flies upon DCV infection that express WD0754 in a wildtype (black) or *Relish^E20^* (red) mutant background. WD0754 transgene expression was acutely induced with 20µM RU (dotted line) compared to non-induced (full line) flies. While induction of WD0754 was protective in a wild type background, it had no beneficial effect upon loss of relish function. (**F**). Survival of female flies upon chronic expression of WD0754 in a homozygous *relish^E20^* mutant background. Expression of WD0754 was induced with 200µM RU486.

To determine if Relish function is essential for the antiviral properties of WD0754, we overexpressed WD0754 in females that carried a *relish* loss-of function mutation and infected these flies with DCV. While acute expression of WD0754 increased the resistance of wildtype flies against DCV, there was no protective effect of WD0754 against DCV when expression was induced in the *relish* mutant background, and these flies even showed a slightly increased susceptibility towards the virus (**Fig 7E**). Thus, relish function is essential for WD0754 mediated DCV resistance. Surprisingly, loss-of relish function partially protected flies against chronic overexpression of WD0754, especially in the latter stage (**Fig 7F**), which might indicate that overactivation of IMD signalling contributes to WD0754 toxicity. IMD-specific immunity does not confer protection to all RNA viruses, with Flock House Virus (FHV) refractory to Relish modulation [31]. We further confirmed the Relish-specific nature of WD0754 conferred immunity by infecting WD0754 expressing flies with FHV. As expected, WD0754 did not protect against FHV infection (**Sup Fig 8A**) and titres of FHV also showed no response (**Sup Fig 8B**). In summary, our data suggest that the *Wolbachia*-derived WD0754 protein increased viral resistance of flies by activating IMD/relish signalling.

## Discussion

The intracellular symbiotic bacterium *Wolbachia pipientis* can protect its arthropod host against viral infections and is used as an agent to reduce the ability of mosquitos to transmit viral diseases including Dengue, Zika, and yellow fever. However, the molecular mechanisms by which *Wolbachia* protects its host against viral infections are only poorly understood. In this study, using the fruit fly *Drosophila melanogaster*, we used the sensitised background of the *dilp2–3,5* mutant to identify a *Wolbachia*-derived protein termed WD0754 that is sufficient to activate the immune response of its host and increase survival of flies in response to viral infection. We further show that WD0754 acts via activation of the Relish transcription factor, a key downstream mediator of the NFkB-like IMD signalling pathway and that Relish function is essential for the antiviral effects of WD0754. Thus, our results provide insights about how *Wolbachia* affects host immunity and this understanding might help to develop novel approaches to prevent the spread of vector borne viral diseases.

The evolutionarily conserved NFkB-like IMD signalling pathway plays an essential role in fly and mosquito immunity [41]. Our data suggest that WD0754 regulates fly immunity via activation of Relish, a key downstream effector of IMD signalling. Upon activation of the IMD pathway, Relish is cleaved and translocates to the nucleus, where it transcribes genes involved in innate immunity, typically targeting gram-negative bacteria [42] [43] but also viruses [44], [45]. WD0754 induction caused both cleavage and nuclear accumulation of Relish, consistent with the hypothesis that it acts via modulation of Relish activity. Furthermore, proteins known to be regulated by Relish were enriched upon WD0754 expression, including Attacin B and C and PGRP-SB1 [40] [36]. We also saw downregulation of proteins which are repressed by IMD activation including Idgf1 [46], consistent with an WD0754-dependent activation of the IMD pathway. Furthermore, CG32368 was the second most highly upregulated protein in the head at 24 hours. This protein is robustly upregulated in response to Sindbis virus [30], DCV [31], Kallithea virus [33] and Zika virus [32]. Expression of CG32368 is increased significantly when IMD repressor *diedel* is absent [30] and is completely blocked when IMD pathway components *dIKKβ* and *dSTING* are not functional [31], again indicating WD0754 exerts anti-viral effects in an IMD dependent manner.

The cytokine Vago was the most up-regulated protein in the head after 24 hours of WD0754 induction. Vago expression is strongly induced upon DCV infection and this expression depends on the function of the DExD/H-box helicase Dicer-2 [47]. Whether, Vago is regulated by Relish in *Drosophila* is currently unclear [47], but in the mosquito *Culex quinquefasciatus* Vago is activated by Relish and activates Jak/STAT signalling [39]. The Jak/STAT pathway is a central component of anti-viral immunity in flies and mosquitoes [45] [48]. Upon viral infection, Jak/STAT signalling activates Vir-1 and Vir-1 is a classical readout for antiviral immunity [49]. Interestingly, Vir-1 was also strongly induced by WD0754 expression, suggesting upregulation of Jak/STAT signalling in response to WD0754 expression. Consistently, other proteins known to be under control of Jak/STAT signalling including Idgf1 and Domeless, the Jak/STAT receptor, were also regulated in response to WD0754 expression. Whether Jak/STAT signalling contributes to the antiviral effects of WD0754 and how exactly WD0754 is able to induce IMD and Jak/STAT signalling is currently unknown and should be addressed in future experiments.

While acute low-level induction of WD0754 was beneficial for immune function, chronic high-level expression was very toxic. Consistently, a prior study also reported embryonic lethality upon constitutive expression of WD0754 [50]. Excessive activation of the IMD pathway may contribute to the toxicity of WD0754. A constitutively active form of the IMD allele expressed ubiquitously results in early pupal lethality [51], demonstrating that chronic activation of IMD signalling can be toxic. Loss-of relish function partially protected flies against WD0754 toxicity, consistent with the hypothesis that excessive activation of IMD signalling contributes to toxicity. Notably, loss of relish function did not fully block WD0754-dependent toxicity, suggesting that WD0754 causes toxicity also via other mechanisms apart from activating IMD signalling. Chronic high-level overexpression of WD0754 also reduced feeding and fecundity. Control of both feeding and egg-laying are well-established adaptive behavioural immune strategies which aim to mitigate the ill-effects of infection [52] [53]. As a master coordinator of immunity, IMD also plays a role in behavioural immunity. IMD expression during larval development affects feeding in adult males [54] and NFkB signalling, which encompasses IMD, in neurons reduces egg laying upon bacteria-induced activation [55]. Other proteins strongly regulated by WD0754 expression have been linked to control of infection-related sickness behaviour. Diuretic-hormone 44 (DH44), is a neuropeptide which plays a role in feeding [56] [57], fecundity [58]), infection [59] [60] and insulin secretion [61]. DH44 was strongly downregulated upon WD0754 induction, suggesting it might contribute to WD0754-dependent phenotypes. The strong reduction in translation-related proteins upon WD0754 induction could also be a protective anti-viral response. Virus infection results in a strong reduction in translation in many species, which is part of an immune response aiming to restrict a key part of the viral life cycle [62]. Recent work has suggested that host translation restricts *Wolbachia* levels but whether *Wolbachia* is able to actively manipulate host translation is still elusive [63]. As WD0754 also caused a reduction in food uptake, it is currently unclear whether the reduction in translation is a direct response towards WD0754 expression or an indirect effect due to partial starvation [64].

Our results suggest that WD0754 expression activates the IMD pathway and thereby protects flies against DCV. However, whether Wolbachia confers viral resistance by priming the immune system is highly debated. For example, IMD pathway genes were not upregulated in viral resistant *Drosophila* flies stably infected with *Wolbachia* [65] and IMD pathway function was not required for *Wolbachia* to block dengue infection in *Drosophila* [66]. However, there is also evidence that *Wolbachia* can directly activate immune pathways. The virulent w*MelPop-CLA Wolbachia* strain activates IMD signalling in the mosquito *Aedes aegypti (*[67]; [68]) and in *Drosophila* [69]. Interestingly, the genes induced by w*MelPop* infection in *Drosophila* include PGRP-SB1, AttB, AttC, Mtk and DptA [69], which were also regulated by induction of WD0754. Noteworthy, Vir-1, Ago2 and Vago expression was not changed in w*MelPop* infected flies. In summary, whether *Wolbachia*-dependent priming of the immune system is controversial, the above data show that Wolbachia can induce IMD activity and expression of genes, which are also regulated by WD0754. In this context, it might also be interesting to study in the future whether WD0754 function contributes to the increased toxicity of virulent *Wolbachia s*trains.

It is at present unknown how or why *Wolbachia* increases insulin signalling in *Drosophila*. The bacterium is transmitted only in the female germline and the evolutionary reproductive interests of the bacterium and the female, but not the male, fly might therefore be identical. It would be interesting to know if *Wolbachia* increases insulin signalling in male flies. We have shown that the density of *Wolbachia* is higher in the *dilp2–3,5* mutant female fly tissues and this may give the bacterium an advantage for entering the female germline, in which case there could be evolutionary conflict between *Wolbachia* and its female host, with the bacterium favouring higher insulin signalling than is optimal for the fly. However, this strain of *Wolbachia* does not appear to increase insulin signalling in wild type females. There may be an interaction with nutritional status. Flies in nature, where the symbiosis evolved, are in general in much lower nutritional status than flies in the laboratory, which would suppress their insulin signalling, and in these circumstances it may be to the advantage of *Wolbachia* to increase it.

A limitation of our study is, that we were not able to show that WD0754 is also essential for *Wolbachia* to protect its host from viral infections. This is due to current technical limitations whereby it is not possible to directly mutate the endogenous WD0754 *Wolbachia* gene. Noteworthy, induction of WD0754 protected flies against DCV but not FHV infection, consistent with the specificity of the IMD pathway [31], while *Wolbachia* has been shown to be protective against both DCV and FHV infection [4]. This finding suggests that additional mechanisms exist apart from WD0754 induction by which *Wolbachia* affects host immunity to provide a broad-spectrum protection against pathogens.

In *Drosophila*, *Wolbachia* has been shown to increase viral resistance of wild type flies [4]. Surprisingly, we did not detect significant differences in DCV resistance in our *wDahomey* wild type strain between flies with and without *Wolbachia*. The reason for this is currently unclear, and might be due to the genetic background of these flies or due to the relative low sensitivity of the survival assay, given that we observed a trend for decreased DCV titre in *Wolbachia* positive wild type flies. Nevertheless, in wDah; *dilp2-3,5* mutants, which had a much higher *Wolbachia* content than wild type flies, there was a clear effect of *Wolbachia* on DCV resistance, indicating that the Wolbachia strain inherent to the Dahomey strain is capable of blocking viral infections. Why *dilp2-3,5* mutants have a higher *Wolbachia* titre is currently unclear. Insulin signalling plays a role in regulation of *Wolbachia* titre in ovaries in response to a high yeast diet and down regulation of TORC1 via rapamycin increased *Wolbachia* titres in ovaries [70]. However, it is also possible that the longer development time of *dilp2-3,5* mutants [23] contributes to the higher *Wolbachia* levels in adult flies, whereby the fly cells are delayed in development while *Wolbachia* keeps proliferating. InR mutant flies are more susceptible to Flavivirus (WNV-Kun) infection [71] and we observed reduced resistance of *dilp2-3,5* mutants negative for *Wolbachia* compared to wild type flies towards DCV infection in one assay. However, this effect was variable and not observed in an independent experiment. Thus, whether IIS signalling is important to protect flies against DCV infection needs to be investigated in future studies.

WD0754 is predicted to be a secreted protein based on the presence of eukaryotic-like domains (ELD), which are also present in eukaryotic genomes and therefore may act as effector in the host cell [29] [72] [73]. However, currently we have no direct evidence, that WD0754 is indeed secreted by Wolbachia and the strong toxicity observed by direct expression of the protein from the *Drosophila* genome might be due to unphysiologically high levels of WD0754 in the host cell. A better understanding of how WD0754 affects IMD signalling and the identification of fly proteins that directly interact with WD0754 might help in future to construct WD0754 transgenes that show lower toxicity but still induce the immune response, a prerequisite for its use as antiviral agent in vector borne diseases. Finally, it needs to be established whether WD0754 can also increase viral resistance of mosquitoes. The finding that a *Drosophila* derived *Wolbachia* strain can protect mosquitos from viral pathogens [74], suggests that the underlying mechanisms, which might include WD0754 induction, are conserved from flies to mosquitoes.

## Materials and methods

### Fly husbandry

The outbred wild-type strain white Dahomey (*wDah*) was used as control genetic background for all experiments. The Dahomey strain was originally collected in Dahomey, now Benin in 1970 and naturally contains the endosymbiontic bacterium Wolbachia [23]. Wolbachia-free flies (*wDahT*) were generated by treating *wDah* flies with Tetracycline [23]. Unless stated otherwise all mutant alleles and transgenic constructs were backcrossed into the *w^Dah^* and *w^DahT^* backgrounds for at least four generations. In order to maintain Wolbachia status and mitochondrial genotype of the background control strain throughout backcrossing, in the first generation, virgin control females (*w^Dah^ or w^DahT^*) were mated to males carrying the respective mutation/transgenic construct. In subsequent generations, female virgins carrying the mutation/transgenic construct were mated to corresponding control males. *Dilp2-3, 5* mutants were originally backcrossed for 10 generations into both the *w^DahT^* and *w^Dah^* background [23] and backcrossing was repeated for another 4 generations before the start of the current study. Flies carrying the *Relish^E20^* loss of function allele were obtained from the Bloomington Drosophila stock centre (RRID:BDSC_9457) and backcrossed for 6 generations into the *Wolbachia*-free *w^DahT^* background. The backcrossed *Relish^E20^* allele was then separately crossed into the UAS-WD0754 and da-GS background. UAS-GR100 flies were published previously [38]. Adult-specific transgene induction was achieved using the GeneSwitch (GS) system [75] using the following driver lines: daughterless (da)-GS for ubiquitous, elav-GS for pan-neuronal, Lsp2-GS for fat body, 5966-GS for enterocyte and Act88f-GS for muscle-specific expression. Transgene induction was achieved by feeding flies that carried both the GS driver line and UAS-construct with RU-486 (RU). RU was dissolved in ethanol and non-induced control flies had the same genotype as induced flies, but received only food with EtOH. *Wolbachia* status was confirmed by PCR prior to experiments using primers directed against the Wolbachia surface protein [26] (**Table 1**). Flies were maintained at 25°C, 65% humidity on a 12hr light 12hr dark cycle and were fed a Sugar Yeast Agar (SYA) diet consisting of sugar (50g/l), yeast (100g/l), agar (1,5g/l), propionic acid and nipagin [76]. Female flies were used for all experiments. Flies were raised at controlled larval densities. Once eclosed, flies were allowed to mate for 48 hours before being sorted under light CO_2_ anaesthesia into experimental vials of 20 individuals.

**Table 1.**
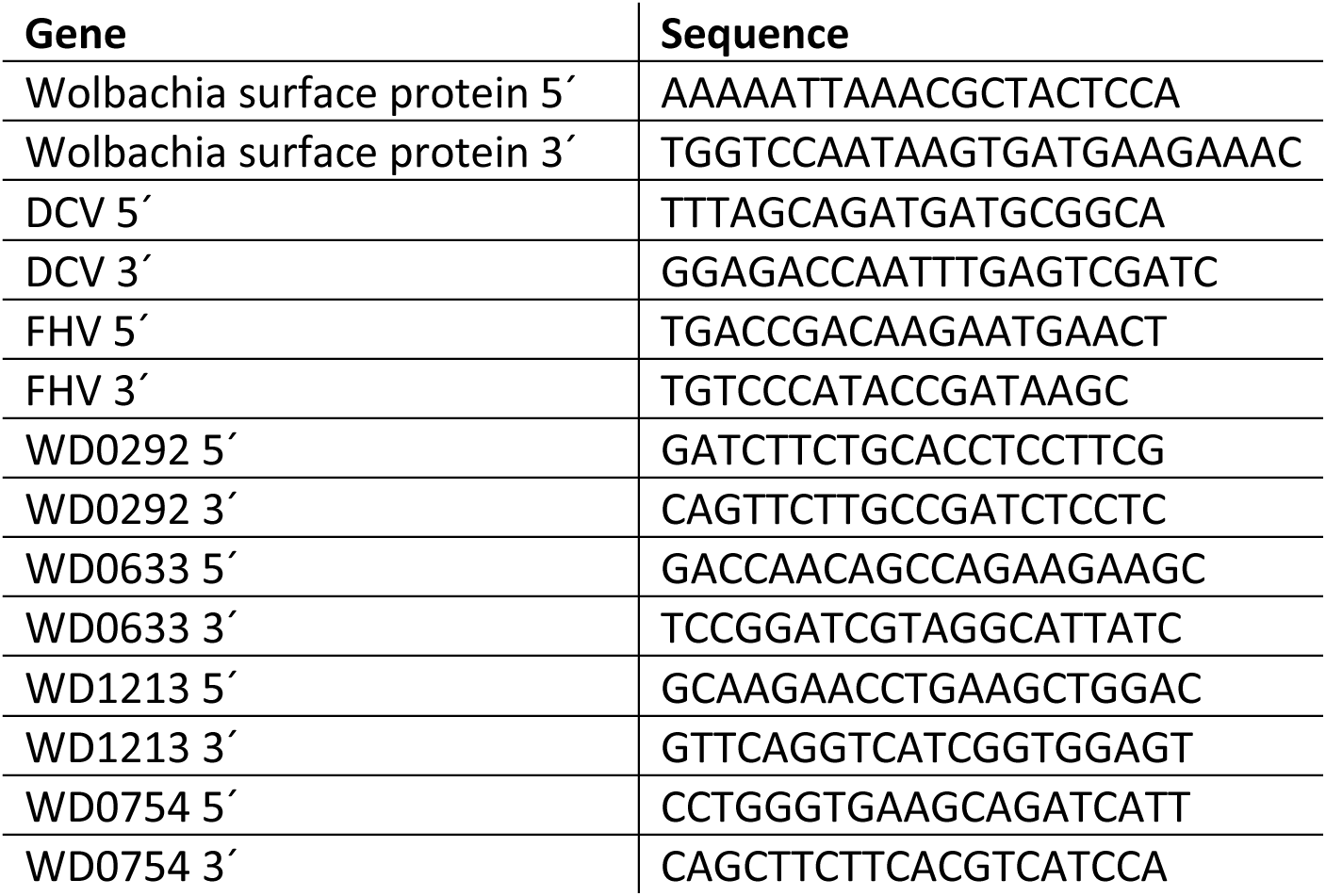
Primers used in this study.

### Generation of transgenic fly lines to express *Wolbachia* ankyrin-domain containing proteins

In order to efficiently express the *Wolbachia* ankyrin-containing proteins WD0754, WD0292, WD1213 and WD0633 in flies, the sequence of the corresponding ORFs was codon optimised matching the *Drosophila melanogaster* codon preference. Codon optimised genes were subsequently synthesised (GenScript Biotech) and cloned into the pUASt attB vector (RRID:Addgene_18944), using KpnI and EcoRI for WD0754, WD0292 and WD1213 and KpnI and NotI for WD0633. Constructs were inserted into the fly genome by φC31-mediated recombination into the attP40 landing site [77].

### Verification of overexpression fly lines by RT-qPCR

To extract total RNA from whole flies, flies were homogenised in Trizol, RNA was phase separated using chloroform and precipitated with sodium acetate and isopropanol and the pellet was then washed with 70% ethanol. Genomic DNA contamination was removed using the TURBO DNAse kit (Thermo Fisher Scientific). cDNA was generated using the SuperScript VILO MasterMix kit with random hexamer primers. RT-qPCR was performed using the Power SYBR Green PCR Master Mix and the primers indicated in **Table 1**. The RT-qPCR was run on a QuantStudio 6 Flex. Results were calculated using the ΔΔCT method and Rpl32 was used for normalization.

### RNA-Seq analysis

Tissue processing, RNA isolation, library preparation, and gene expression analysis for the Drosophila host genes were previously described [26]. In brief, brain, gut, thorax and fat body of 10-day old female wDah, wDahT control and wDah; *dilp2-3,5* and wDahT; *dilp2-3,5* mutant flies were dissected and RNA was extracted using Trizol, according to the manufacturer’s guidelines. Three independent biological replicates were used per genotype. Stranded TruSeq RNA-seq library preparation was then performed on total RNA after rRNA depletion (Ribo-zero). Libraries were sequenced with 37 mio, 100 bp single reads on an Illumina HiSeq2500 (Illumina). Adapter trimming was carried out using flexbar (version 2.5, RRID:SCR_013001), using minimum read length after trimming of 30, and quality threshold of 20. Transcripts were then mapped to the BDGP6.28 reference genome using tophat2 (version 2.1.0, RRID:SCR_013035) and counted via summarizeOverlaps (part of the Bioconductor R package GenomicAlignments, version 1.10.1, RRID:SCR_006442) with the option ‘intersectionNotEmpty’. FPKM were calculated using DESeq2 (version 1.14.1, RRID:SCR_015687).. To determine Wolbachia gene expression, we separately identified Wolbachia associated transcripts using Trinity de-novo and genome-guided (Wolbachia pipientis) assemblies [78]. This was followed by a stringent BLAST search to identify ORFs. Differential expression analysis was carried out tissue-wise using RSEM [79]. Differential expression analysis was carried out tissue-wise using DESeq2, testing the insulin response. p-Values were adjusted for multiplicity by DESeq2 using the Benjamini-Hochberg procedure on a per-tissue basis, with independent filtering enabled.

### Lifespan analysis

Female flies were generated as described above and sorted onto either control food (containing just ethanol) or RU at the indicated dose. Flies were transferred to new food and scored for any deaths every 2-3 days. All lifespan experiments were done using female flies.

### Body weight and feeding assays

Flies were weighed in pairs and body weight averaged. Feeding was measured by uptake of blue dye added to the food [80].

### Peptide preparation for proteomics

Per treatment, six replicates of 10 fly tissues (head, gut or fat) were homogenised using a pestle mounted to a handgun and subsequently lysed in 6M guanidine chloride, 10mM TCEP, 40 mM 2-Chloroacetamide, 100mM Tris pH 8.5 lysis buffer. After shaking at 700rpm and 95°C for 10 minutes, samples were sonicated using the Bioruptor plus for 10 cycles. After spinning down, 300 µg of the supernatant was diluted 10-fold with digestion buffer (20mM Tris 8.5 pH, 10% acetonitrile). 3µg Trypsin were added and samples were digested at 37°C overnight. Peptides were cleaned using StageTips produced in house by the Proteomics Core Facility. The Tips were first cleaned using methanol, 0.1% Formic Acid (FA) in 40% Acetonitrile and 0.1% FA in water. Proteins were then washed two times using 0.1% FA in water. Peptides were then dried using the Speed-Vac, resuspended in 0.1% FA and measured. 5µg per replicate were again dried and given to the Proteomics facility for LC-MS/MS analysis.

### LC-MS/MS analysis

Peptides were separated on a 40 cm, 75 μm internal diameter packed emitter column (Coann emitter from MS Wil, Poroshell EC C18 2.7 micron medium from Agilent) using an EASY-nLC 1200 (Thermo Fisher Scientific). The column was maintained at 50°C. Buffer A and B were 0.1% formic acid in water and 0.1% formic acid in 80% acetonitrile, respectively. Peptides were separated on a segmented gradient from 4% to 31% buffer B for 59 min at 300 nl / min, followed by a higher organic wash. Eluting peptides were analyzed on a Orbitrap Fusion LUMOS Tribrid mass spectrometer equipped with a FAIMS Pro interface (Thermo Fisher Scientific) in Boxcar-DIA mode. Peptide precursor m/z measurements were carried out at 60000 resolution in the 400 to 800 m/z range followed by 2 10 Boxcar tSIM scans and finally 29 DIA scans with an isolation width of 14 Th and a resolution of 15000. All data were acquired at a FAIMS CV of −50 V. These acquisition parameters yielded five quantification points per peak.

### Protein identification, quantification and analysis

Raw protein data were analysed with Spectronaut 16 (Biognosys) using dynamic default parameters against a combined fasta database of 21,939 *Drosophila melanogaster* and 1159 *Wolbachia pipientis wMel* sequences (Uniprot). Methionine oxidation and protein N-terminal acetylation were set as variable modifications; cysteine carbamidomethylation was set as fixed modification. The digestion parameters were set to “specific” and “Trypsin/P,” with two miscleavages permitted. Differential expression analysis was performed using limma, version 3.34.9, [81] in R, version 3.4.3 (R Core Team 2017). Gene ontology enrichment analysis was conducted using ShinyGO (V0.77) [82].

### Viral infections

For WD1213, WD0633 and WD0292 induction of transgene expression was initiated 72h prior to infection by feeding flies with 200µM RU and flies stayed on the RU food throughout the experiment. As the prolonged induction of WD0754 prior to infection negatively interfered with the flies injury response, WD0754 transgene expression was initiated 12h prior infection by transferring flies to food containing 10-20µM RU. WD0754 expressing flies remained on the RU food throughout the experiment. Infections were conducted using a needle prick method [83]. A 0.10mm stainless steel minutien pin was first dipped in sterile PBS and used to infect sham control flies. The same pin was subsequently dipped in a 3.4 x10^9^ PFU DCV or 7.6 x10^6^ PFU FHV solution and pricked in the plural suture in the flies thorax. Flies were scored every 2-3 days until death began and every day thereafter.

### Viral quantification

For quantification of DCV and FHV titres, flies were pooled in groups of 10-20 and flash frozen in liquid nitrogen 24 hours post-infection. RNA was subsequently extracted, cDNA generated and viral titre was measured using RT-qPCR as described above. Primers used to quantify DCV and FHV are listed in **Table 1**.

### Immunofluorescence

Relish localisation in fat body tissue was analyzed as previously described [84]. Therefore, fat body tissues were dissected in PBS and fixed 30 min with 4% formaldehyde, methanol-free (ThermoFisher, #28908). Samples were washed in 0.3% Triton-X/PBS (PBST), blocked in 5% bovine serum albumin (BSA)/PBST for 1 h at room temperature, incubated in primary antibody overnight at 4°C, and in secondary antibody for 2 h at room temperature. The following primary antibodies were used: anti-Relish (Developmental Studies Hybridoma Bank, #21F3, 1:100). The following secondary antibodies were used: Alexa Flour 594 goat anti-mouse IgG (ThermoFisher, #A11005, 1:1,000). Samples were mounted with VECTASHIELD Antifade Mounting Medium with DAPI (Vector Laboratories, #H-1200). Three images per sample were captured using a Leica TCS SP8 DLS confocal microscope with 20x objective and 6x digital zoom. For Relish protein localisation, images were processed by background subtraction and median filtering using Imaris 9 (Bitplane). The number of nuclei with high Relish fluorescence intensity was normalized by the total number of nuclei. Confocal settings were kept consistent between images of the same experiment. Statistical analyses were performed in R 4.1.0. Linear mixed model was generated and analysed in R using lme4, lmertest, and emmeans package.

### Immunoblotting

Whole flies were homogenized and lysed in ice-cold RIPA buffer supplemented with cOmplete, Mini, EDTA-free, Protease Inhibitor Cocktail (Roche) and PhosSTOP phosphatase inhibitor tablet (Roche) using a hand-held homogenizer. Extracts were centrifuged and protein concentrations were determined using Pierce BCA Protein Assay Kit (ThermoFisher). Extracts were mixed with 4x Laemmli loading buffer and boiled for 5 min at 95°C. 10 µg of protein per lane were separated on Any kD Criterion TGX stain-free precast gels (Bio-Rad) and transferred to Immobilon-FL PVDF membranes (Millipore). Membranes were blocked by Intercept TBS Blocking Buffer (LI-COR) for 1 h and probed with the following primary antibodies diluted in Intercept T20 TBS Antibody Diluent (LI-COR): anti-Relish (Developmental Studies Hybridoma Bank, #21F3, 1:100). The following secondary antibodies were used: IRDye 800CW Goat anti-Mouse IgG (H + L) (LI-COR, #926-32210, 1:15,000). Total protein on the membrane was visualized as stain-free signal using ChemiDoc MP Imagers (Bio-Rad) and was used for normalizing protein expression levels. Immunoblotting images were captured using Odyssey Infrared Imaging system with application software V3.0.30 (LI-COR) and were analysed using Fiji^109^ (US National Institutes of Health).

## Acknowledgments

We would like to thank Prof Jean-Luc Imler and the members of his lab for kindly sharing DCV and FHV virus samples with us. We acknowledge Xinping Li, Thomas Colby and Ilian Atanassov from the Proteomics Core Facility at the Max Planck Institute for Biology of Ageing for generating the proteomics data. Christian Kukat and the FACS and Imaging Core Facility at the Max Planck Institute for Biology of Ageing are acknowledged for their support with confocal microscopy. We thank Robert Sehlke for bioinformatic assistance. Emilie Funk, Jenny Fröhlich, Jacqueline Esser and Andre Pahl are acknowledged for technical assistance. Stocks obtained from the Bloomington Drosophila Stock Center (NIH P40OD018537) were used in this study.

## Author Contributions

**TL**: Conceptualization, Writing – Original Draft Preparation, Writing – Review & Editing, Formal Analysis, Supervision, Investigation, Project Administration, Methodology, Visualization

**BT**: Investigation, Formal Analysis, Project Administration, Visualization

**RLM**: Investigation

**LST**: Investigation

**PZ**: Investigation, Formal Analysis, Visualization

**SG**: Conceptualization, Writing – Review & Editing, Formal Analysis, Supervision, Investigation, Methodology, Visualization

**LP**: Funding Acquisition, Supervision, Conceptualization, Methodology, Writing – Review & Editing

## Supporting information

Supplemental Figures 1-8.

**Figure S1.**
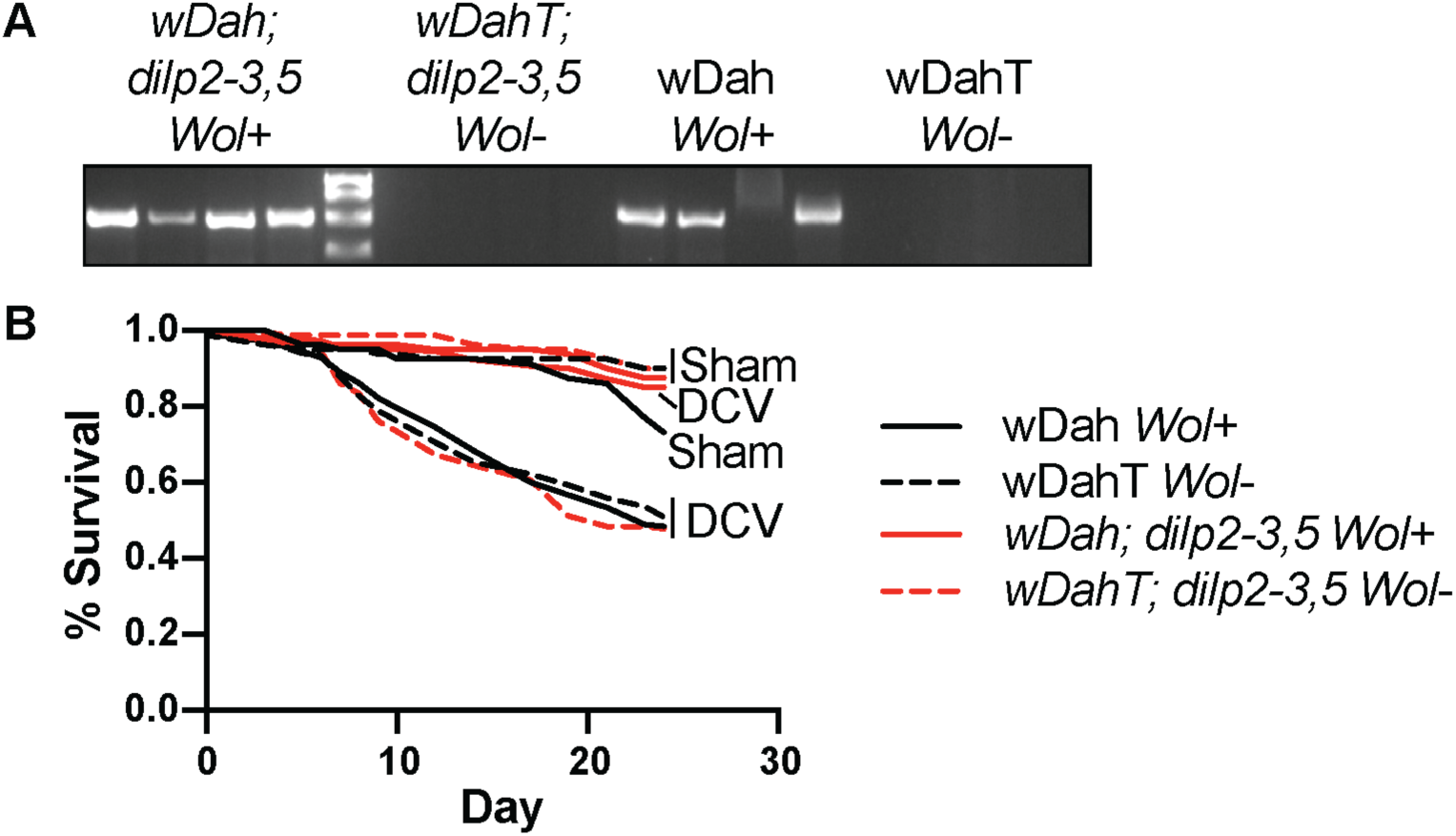
DCV resistance of *dilp2,3-5* mutants is dependent on *Wolbachia*. (**A**) PCR verification of *Wolbachia* status of *dilp2-3,5* and wt flies using primers against the Wolbachia surface protein. Each lane represents a single fly. A band indicates that the fly carried *Wolbachia* (*Wol*). (**B**) Survival of *dilp2-3,5* and wildtype (wt) females, with (Wol+, full line) and without (Wol-, dotted line) *Wolbachia* upon infection with DCV. *dilp2-3,5 Wolbachia*-positive flies were significantly more resistant towards DCV infection than *dilp2-3,5* flies without *Wolbachia* and wt control flies (****: p < 0.0001, Gehan-Breslow-Wilcoxon test, n = 100 for sham and n = 180 for DCV infected flies. Independent experiment of Figure 1A.

**Figure S2.**
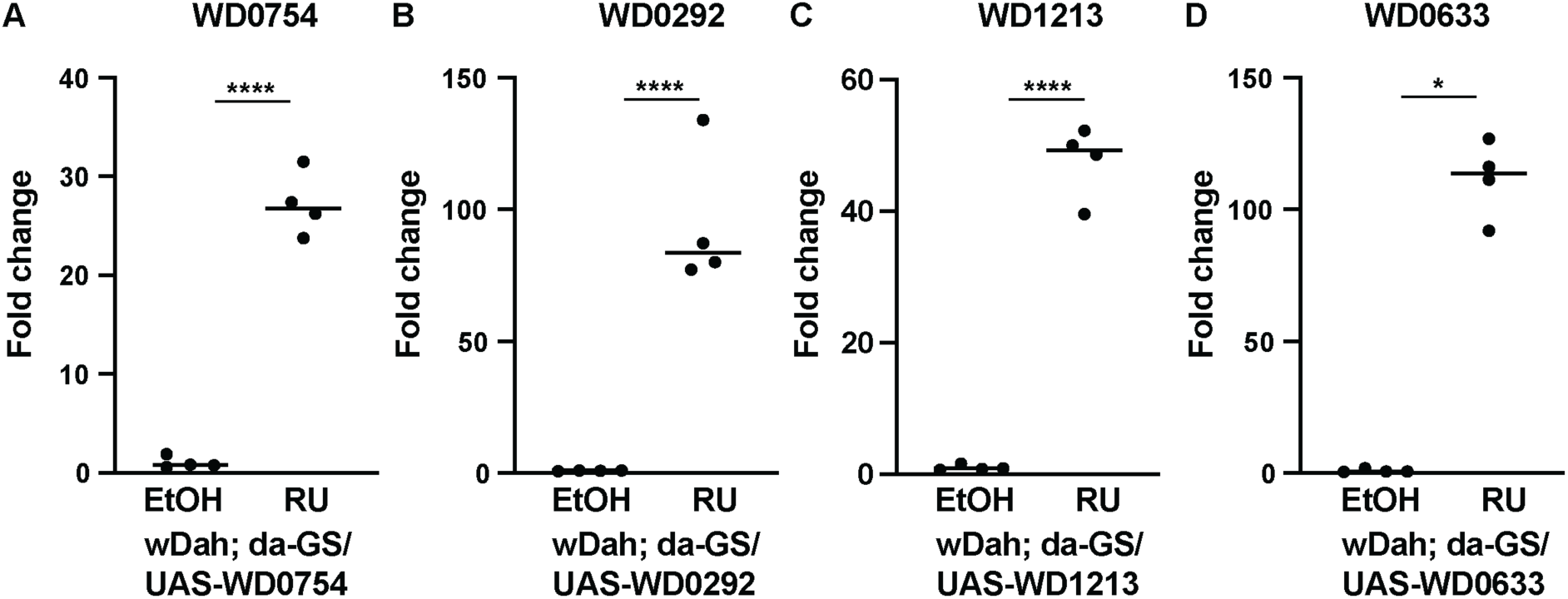
RT-qPCR verification of *Wolbachia* ankyrin transgene expression. RT-qPCR analysis of whole flies expressing the (**A**) WD0754, (**B**) WD0292, (**C**) WD1213 and (**D**) WD0633 ankyrin-domain containing *Wolbachia* constructs under the control of the ubiquitous da-GS driver line. Expression was induced with 200µM RU486 (induced) and compared to control (non-induced, EtOH only) flies. All constructs were significantly expressed. Note that due to the codon optimization of the transgenic constructs the probes used for the RT-qPCR analysis do not detect the endogenous *Wolbachia* transcripts, i.e. the fold change does not indicate the levels relative to the endogenous level of the respective ankyrin domain containing transcript. *p<0.05, ****p<0.0001, Mann-Whitney test. n = 4 replicates with 20 flies each.

**Figure S3.**
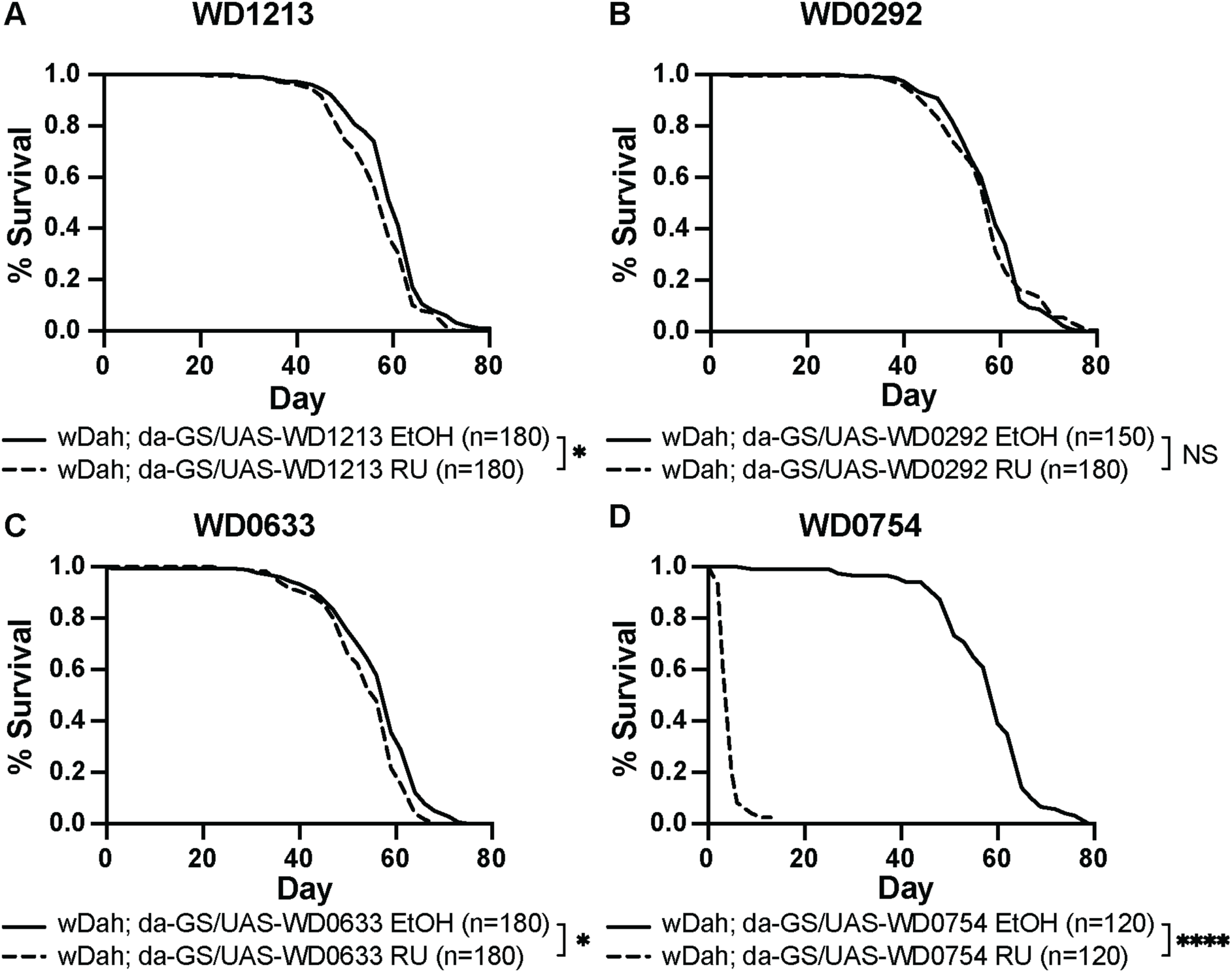
Overexpression of WD0754 shortens lifespan in a *Wolbachia* positive background. ***(A-D)*** Survival of female flies ubiquitously overexpressing the ankyrin-domain containing *Wolbachia* proteins (**A**) WD1213, (**B**) WD0292, (**C**) WD0633, (**D**) and WD0754 under the control of the da-GS driver. Expression of the transgene was induced with 200µM RU486 (induced). Overexpression of WD1213, WD0292 and WD0633 had only minor or no effect on fly survival. In contrast, overexpression of WD0754 drastically shortened lifespan (*: p < 0.05, ****: p < 0.0001, log-rank test, n=180 females per treatment for A-C except for 0292 control: n=150, n=120 females per treatment for d). All flies contained *Wolbachia*, in contrast to Figure 3, which shows the corresponding data for *Wolbachia*-negative flies.

**Figure S4.**
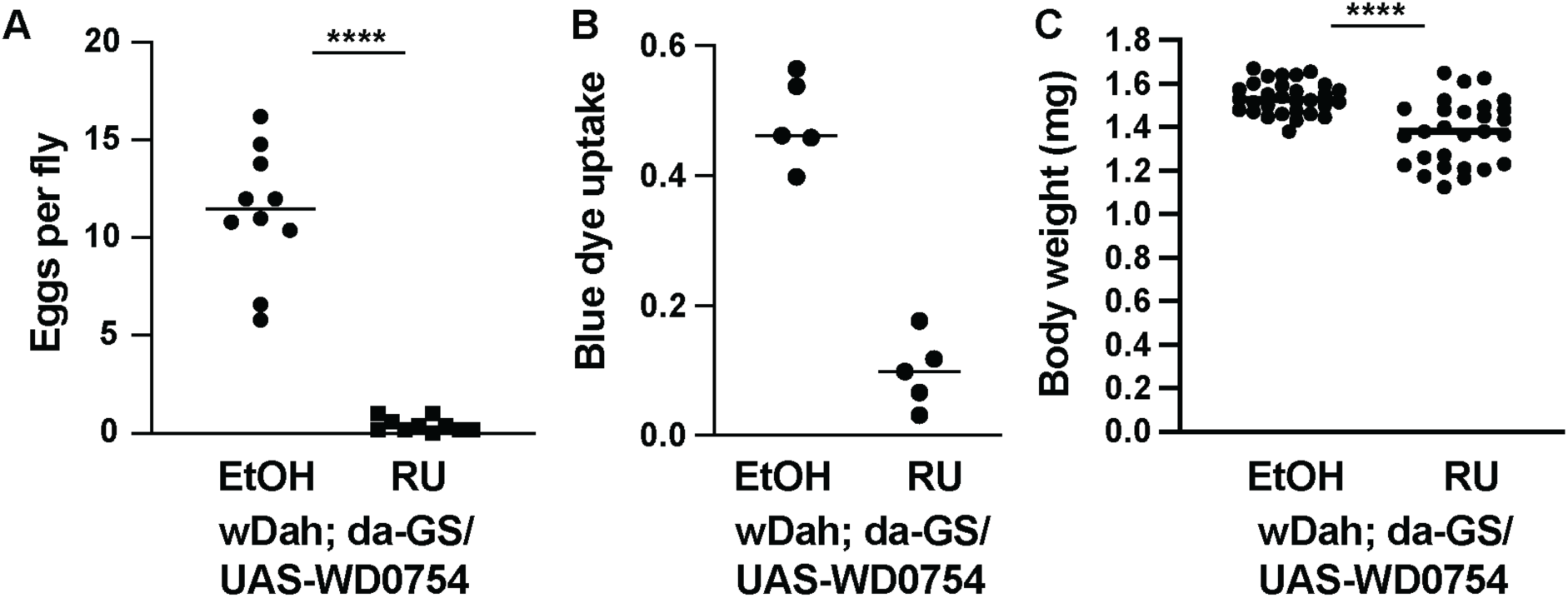
Chronic expression of WD0754 reduced fecundity, food uptake and body weight. Ubiquitous expression of WD0754 under the control of the da-GS driver line reduced. (**A**) egg laying, (**B**) food uptake and (**C**) body weight. Expression was induced using 50µM RU. (**A**) Egg laying was measured by counting the number of eggs laid by 10 groups of 5 flies over a period of 15 h. (**B**) Food uptake was measured by allowing flies to feed on blue food for 45 minutes before they were homogenized and the absorbance was measured, n = 5 replicates of 10 pooled flies. (C) For the determination of body weight, female flies were weighed in pairs and then averaged, n = 28-30 pairs. ****p<0.0001, Mann-Whitney test.

**Figure S5.**
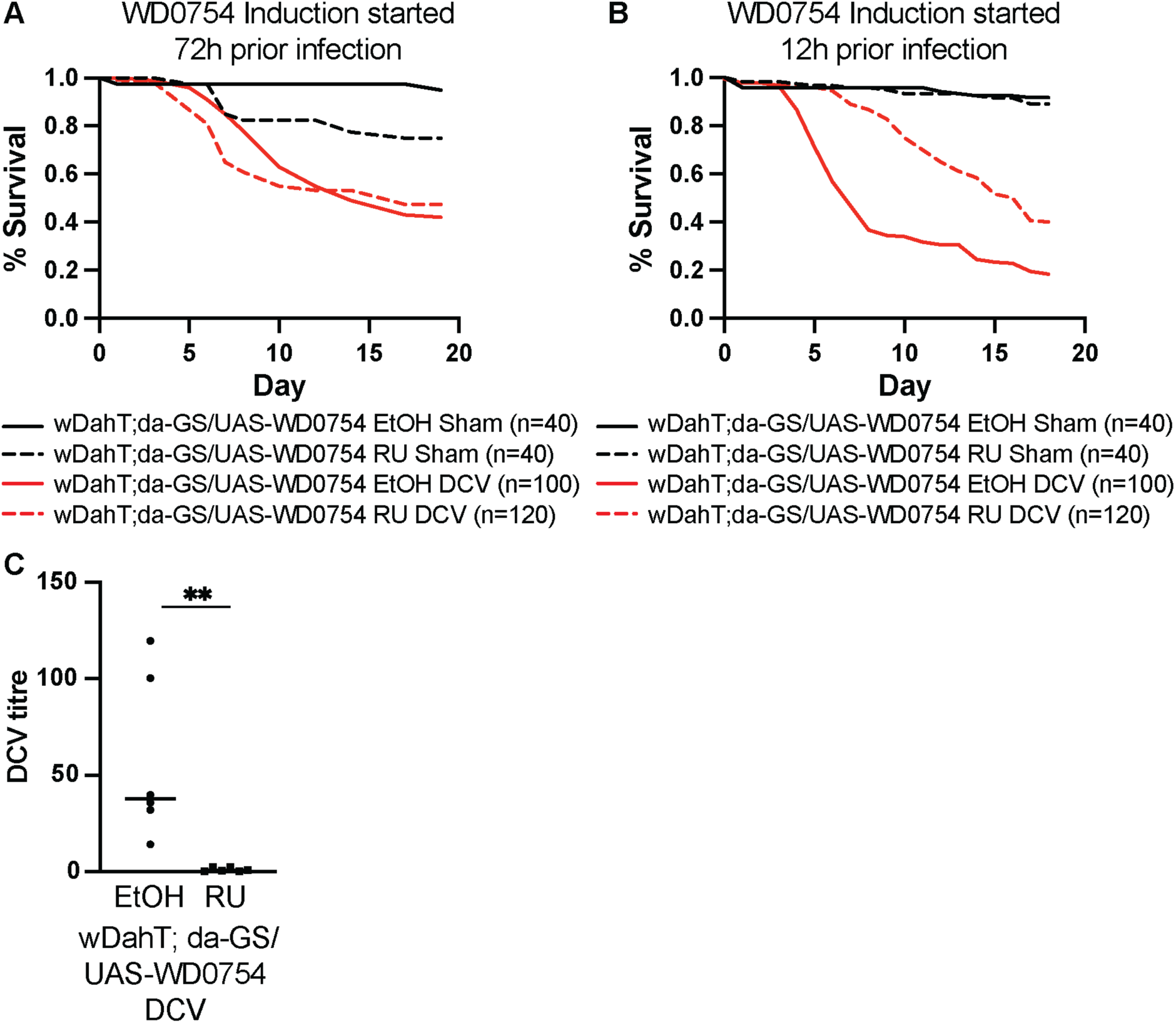
Acute but not chronic induction of WD0754 protects flies against DCV. (**A**) Survival of female flies that ubiquitously expressed WD0754 under the control of the da-GS driver in response to DCV infection. Expression was induced using 10 µM RU486 three days before flies were infected with DCV. Chronic expression of WD0754 did not protect flies against DCV infection. Sham infection caused an increase in mortality in flies that overexpressed WD0754, but not in sham control animals (n = 40 for sham, n = 100 for WD0754 + DCV and n = 120 for Induced + DCV). (**B**) Acute induction of WD0754 with 10 µM RU486 12 hours prior to infection did significantly increase the viral resistance of female flies (p<0.0001 for WD0754 + DCV vs Induced + DCV, Gehan-Breslow-Wilcoxon test, n = 40 for sham, n = 100 for WD0754 + DCV and n = 120 for Induced + DCV). **(C**) RT-qPCR measurement of DCV titre upon acute induction of WD0754 48h post infection. Data are shown as fold change relative to the induced condition. **p<0.005, Mann Whitney test n = 6 groups of 20 pooled females.

**Figure S6.**
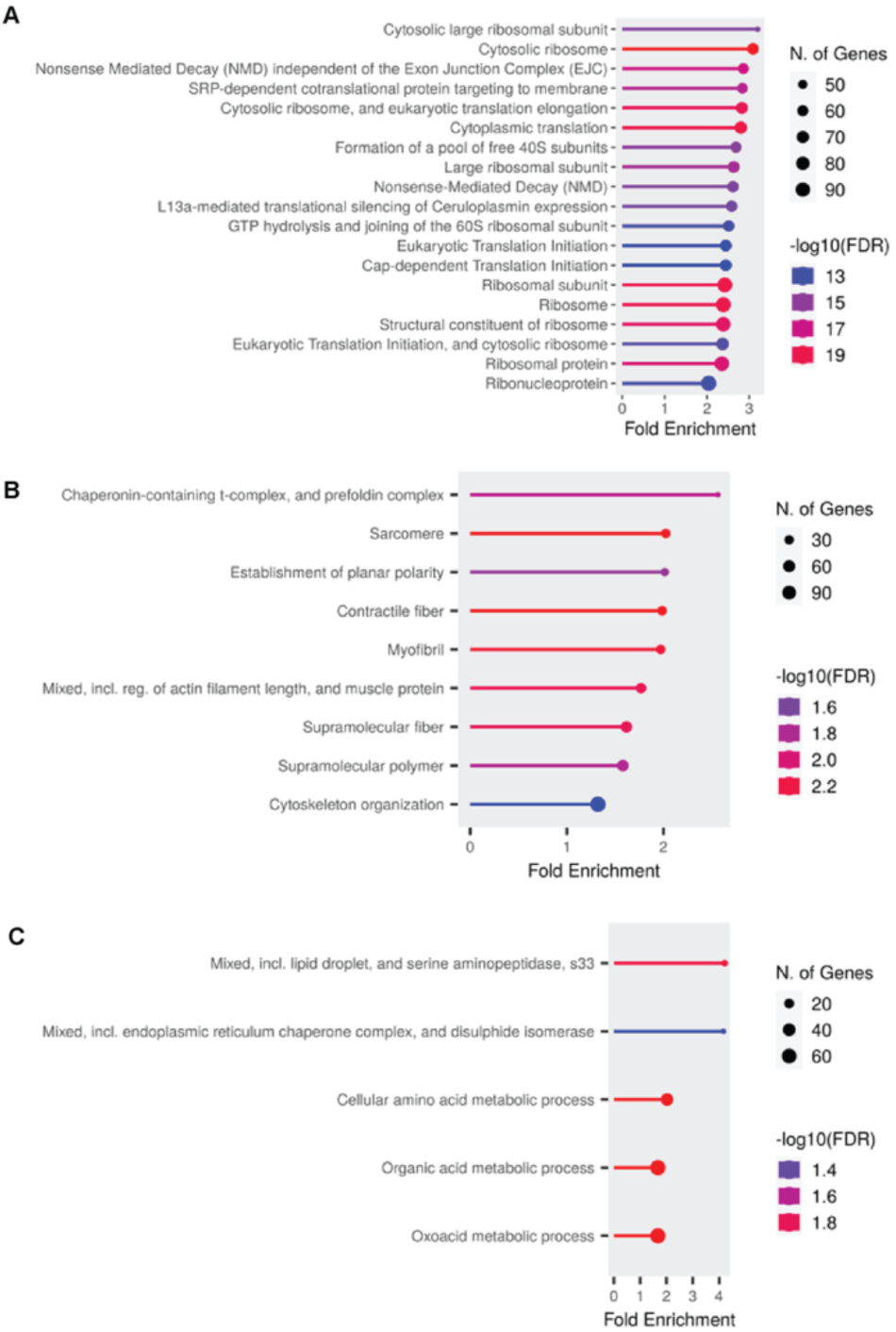
Gene ontology enrichment after 72 hours of WD0754 expression. Gene ontology enrichment analysis of proteins that were significantly different between female WD0754 expressing flies and controls in the head (**A**), gut (**B**) and fat body (**C**) 72 hours after induction of the transgene. Flies were positive for *Wolbachia*.

**Figure S7.**
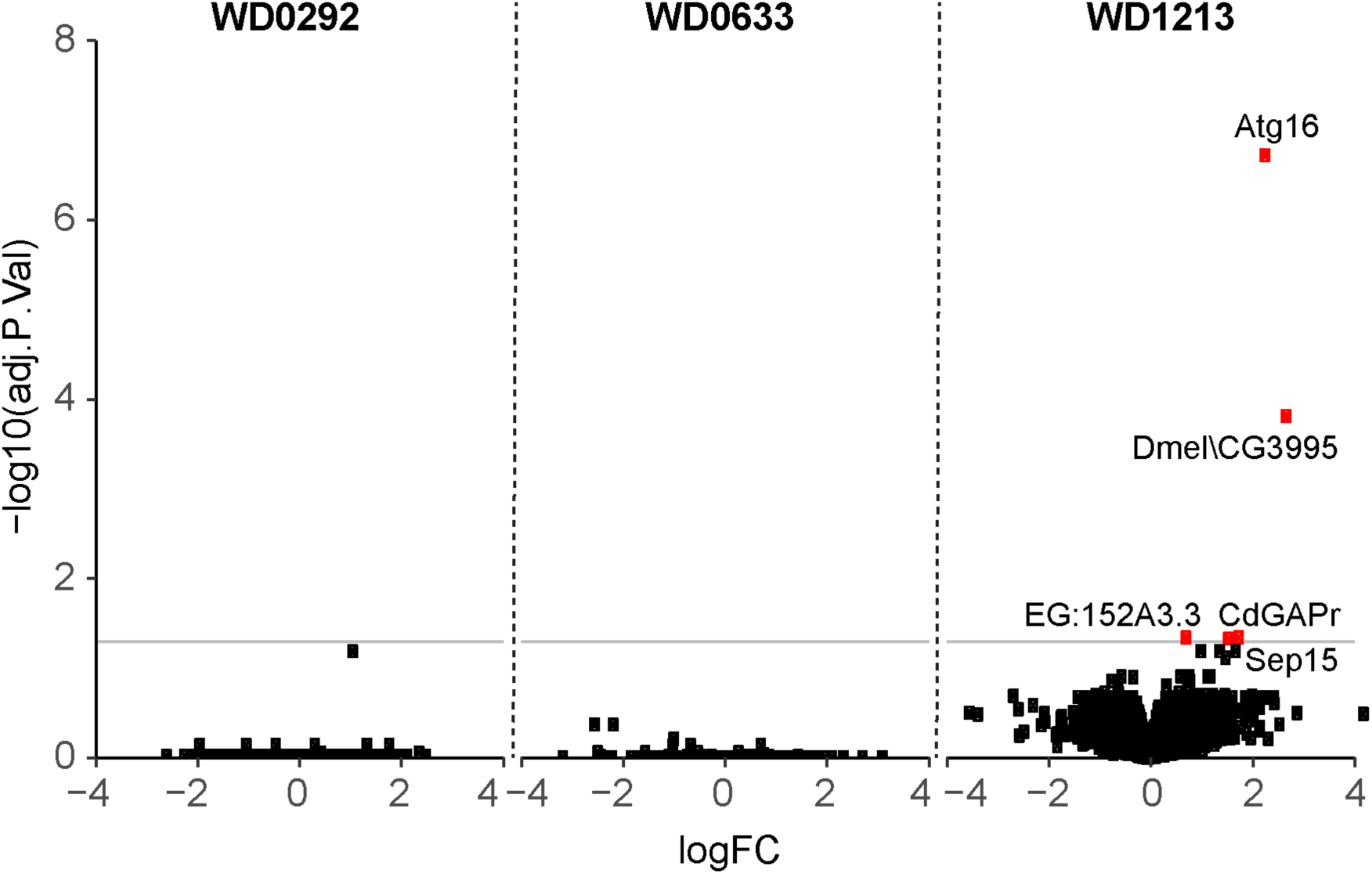
Overexpression of WD0292, WD0633 and WD1213 does not strongly affect the fly head proteome. Changes in the head proteome upon overexpression of WD0292, WD0633 and WD1213 for 72h were measured by mass spectrometry. Transgene expression was induced with 200µM RU486 using the ubiquitous *da*-GS driver line. The Red dots indicate proteins significantly regulated upon WD1213 expression (adjusted P < 0.05, n=6 biological replicates containing 10 heads each).

**Figure S8.**
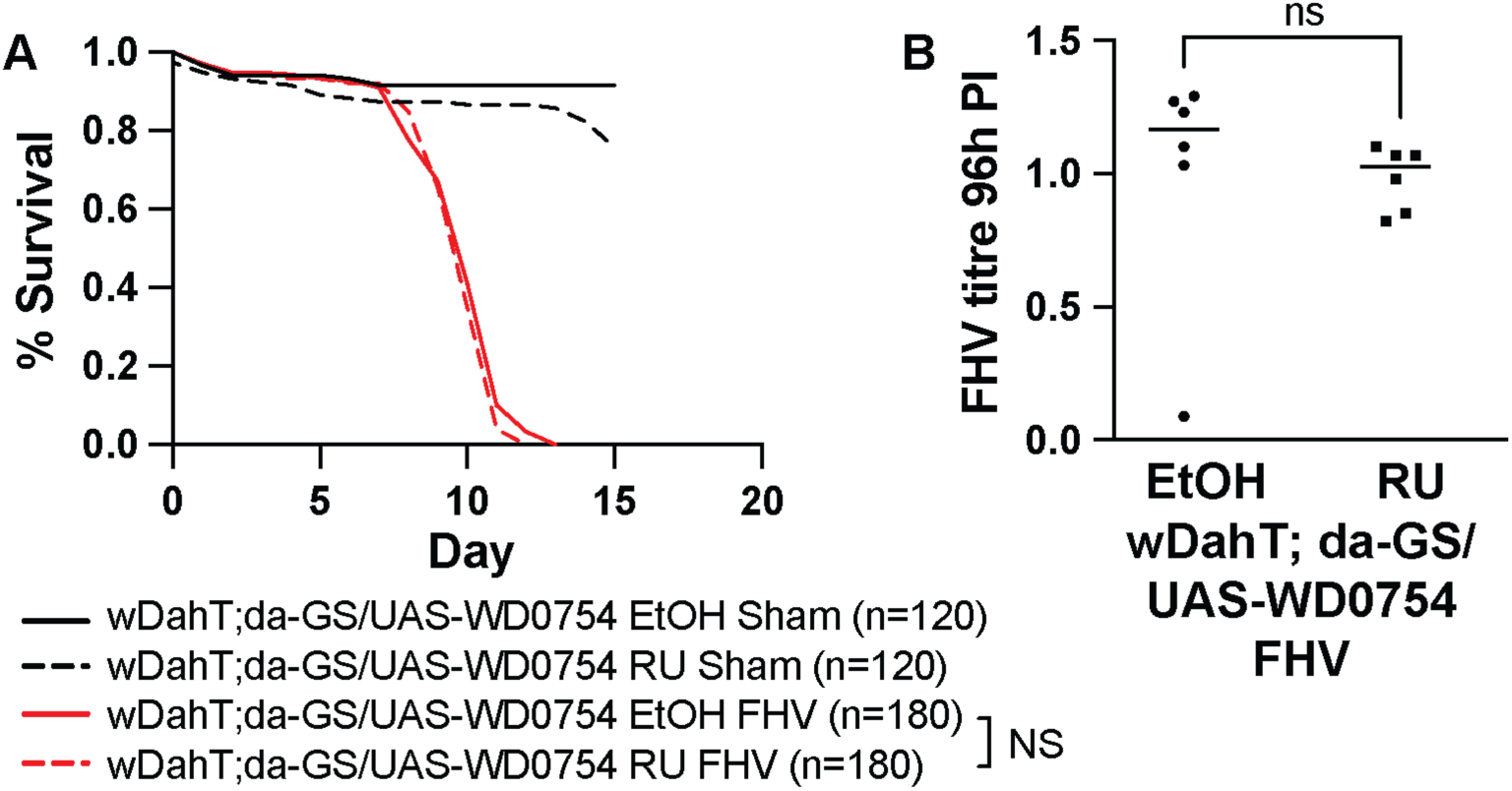
Acute induction of WD0754 did not protect against Flock House Virus (FHV). (**A**) Survival of female flies upon infection with FHV. Acute induction of WD0754 with 10 µM RU486 12 hours prior to infection with FHV did not increase the viral resistance of flies. (n = 120 for sham and n = 180 for FHV infected flies). (**B**) RT-qPCR measurement of FHV titre in flies with acute expression of WD0754. FHV titre was measured 96 hours post infection (PI) and levels are shown relative to the induced condition. WD0754 induction did not significantly decrease FHV levels (Mann-Whitney test, n = 6 replicates of 20 pooled females).

## Reference List

1. Werren JH, Baldo L, Clark ME. Wolbachia: master manipulators of invertebrate biology. Nature Reviews Microbiology. 2008;6:741. doi: 10.1038/nrmicro1969.

2. Werren JH, Zhang W, Guo LR. Evolution and phylogeny of Wolbachia: reproductive parasites of arthropods. Proceedings of the Royal Society B (Biological Sciences). 1995;261(1360):55–63. Epub 1995/07/22. doi: 10.1098/rspb.1995.0117. PubMed PMID: 7644549.

3. Hedges LM, Brownlie JC, O’Neill SL, Johnson KN. Wolbachia and virus protection in insects. Science. 2008;322(5902):702-. doi: 10.1126/science.1162418.

4. Teixeira L, Ferreira Á, Ashburner M. The bacterial symbiont Wolbachia induces resistance to RNA Viral infections in Drosophila melanogaster. PLOS Biology. 2008;6(12):e1000002. doi: 10.1371/journal.pbio.1000002.

5. Mains JW, Kelly PH, Dobson KL, Petrie WD, Dobson SL. Localized control of Aedes aegypti in Miami, FL, via inundative releases of Wolbachia-infected male mosquitoes. Journal of Medical Entomology. 2019;56(5):1296–303. doi: 10.1093/jme/tjz051.

6. Utarini A, Indriani C, Ahmad RA, Tantowijoyo W, Arguni E, Ansari MR, et al. Efficacy of Wolbachia-infected mosquito deployments for the control of Dengue. New England Journal of Medicine. 2021;384(23):2177–86. doi: 10.1056/NEJMoa2030243. PubMed PMID: 34107180.

7. Lindsey ARI, Bhattacharya T, Newton ILG, Hardy RW. Conflict in the intracellular lives of endosymbionts and viruses: A mechanistic look at Wolbachia-mediated pathogen-blocking. Viruses. 2018;10(4). Epub 20180321. doi: 10.3390/v10040141. PubMed PMID: 29561780; PubMed Central PMCID: PMCPMC5923435.

8. White PM, Serbus LR, Debec A, Codina A, Bray W, Guichet A, et al. Reliance of Wolbachia on high rates of host proteolysis revealed by a genome-wide RNAi screen of Drosophila cells. Genetics. 2017;205(4):1473–88. Epub 20170203. doi: 10.1534/genetics.116.198903. PubMed PMID: 28159754; PubMed Central PMCID: PMCPMC5378107.

9. Ferree PM, Frydman HM, Li JM, Cao J, Wieschaus E, Sullivan W. Wolbachia utilizes host microtubules and Dynein for anterior localization in the Drosophila oocyte. PLOS Pathogens. 2005;1(2):e14. Epub 20051014. doi: 10.1371/journal.ppat.0010014. PubMed PMID: 16228015; PubMed Central PMCID: PMCPMC1253842.

10. Newton IL, Savytskyy O, Sheehan KB. Wolbachia utilize host actin for efficient maternal transmission in Drosophila melanogaster. PLOS Pathogens. 2015;11(4):e1004798. Epub 20150423. doi: 10.1371/journal.ppat.1004798. PubMed PMID: 25906062; PubMed Central PMCID: PMCPMC4408098.

11. Sheehan KB, Martin M, Lesser CF, Isberg RR, Newton IL. Identification and characterization of a candidate Wolbachia pipientis Type IV effector that interacts with the actin cytoskeleton. mBio. 2016;7(4). Epub 20160705. doi: 10.1128/mBio.00622-16. PubMed PMID: 27381293; PubMed Central PMCID: PMCPMC4958246.

12. Caragata EP, Rances E, Hedges LM, Gofton AW, Johnson KN, O’Neill SL, et al. Dietary cholesterol modulates pathogen blocking by Wolbachia. PLOS Pathogens. 2013;9(6):e1003459. Epub 20130627. doi: 10.1371/journal.ppat.1003459. PubMed PMID: 23825950; PubMed Central PMCID: PMCPMC3694857.

13. Molloy JC, Sommer U, Viant MR, Sinkins SP. Wolbachia modulates lipid metabolism in Aedes albopictus mosquito cells. Applied Environmental Microbiology. 2016;82(10):3109–20. Epub 20160502. doi: 10.1128/AEM.00275-16. PubMed PMID: 26994075; PubMed Central PMCID: PMCPMC4959074.

14. Geoghegan V, Stainton K, Rainey SM, Ant TH, Dowle AA, Larson T, et al. Perturbed cholesterol and vesicular trafficking associated with dengue blocking in Wolbachia-infected Aedes aegypti cells. Nature Communications. 2017;8(1):526. Epub 20170913. doi: 10.1038/s41467-017-00610-8. PubMed PMID: 28904344; PubMed Central PMCID: PMCPMC5597582.

15. Wong ZS, Brownlie JC, Johnson KN. Oxidative stress correlates with Wolbachia-mediated antiviral protection in Wolbachia-Drosophila associations. Applied and Environmental Microbiology. 2015;81(9):3001–5. doi: doi:10.1128/AEM.03847-14.

16. Hussain M, Zhang G, Leitner M, Hedges LM, Asgari S. Wolbachia RNase HI contributes to virus blocking in the mosquito Aedes aegypti. iScience. 2023;26(1):105836. Epub 2023/01/14. doi: 10.1016/j.isci.2022.105836. PubMed PMID: 36636344; PubMed Central PMCID: PMCPMC9830209.

17. Gupta V, Vasanthakrishnan RB, Siva-Jothy J, Monteith KM, Brown SP, Vale PF. The route of infection determines Wolbachia antibacterial protection in Drosophila. Proc Biol Sci. 2017;284(1856). Epub 2017/06/09. doi: 10.1098/rspb.2017.0809. PubMed PMID: 28592678; PubMed Central PMCID: PMCPMC5474083.

18. Perlmutter JI, Schedl ME, Unckless RL. Wolbachia enhances the survival of Drosophila infected with fungal pathogens. bioRxiv. 2023:2023.09.30.560320. doi: 10.1101/2023.09.30.560320.

19. Partridge L, Alic N, Bjedov I, Piper MDW. Ageing in Drosophila: The role of the insulin/Igf and TOR signalling network. Experimental Gerontology. 2011;46(5):376–81. doi: 10.1016/j.exger.2010.09.003.

20. Becker T, Loch G, Beyer M, Zinke I, Aschenbrenner AC, Carrera P, et al. FOXO-dependent regulation of innate immune homeostasis. Nature. 2010;463(7279):369–73. doi: 10.1038/nature08698. PubMed PMID: 20090753.

21. Haqshenas G, Terradas G, Paradkar PN, Duchemin JB, McGraw EA, Doerig C. A Role for the Insulin Receptor in the Suppression of Dengue Virus and Zika Virus in Wolbachia-Infected Mosquito Cells. Cell Rep. 2019;26(3):529–35 e3. doi: 10.1016/j.celrep.2018.12.068. PubMed PMID: 30650347.

22. Ikeya T, Broughton S, Alic N, Grandison R, Partridge L. The endosymbiont Wolbachia increases insulin/IGF-like signalling in Drosophila. Proceedings of the Royal Society B (Biological Sciences). 2009;276(1674):3799–807. Epub 2009/08/19. doi: 10.1098/rspb.2009.0778. PubMed PMID: 19692410.

23. Grönke S, Clarke D-F, Broughton S, Andrews TD, Partridge L. Molecular evolution and functional characterization of Drosophila insulin-like peptides. PLOS Genetics. 2010;6(2):e1000857-e. doi: 10.1371/journal.pgen.1000857. PubMed PMID: 20195512.

24. Osborne SE, Leong YS, O’Neill SL, Johnson KN. Variation in antiviral protection mediated by different Wolbachia strains in Drosophila simulans. PLOS Pathogens. 2009;5(11):e1000656. doi: 10.1371/journal.ppat.1000656.

25. Martinez J, Longdon B, Bauer S, Chan Y-S, Miller WJ, Bourtzis K, et al. Symbionts commonly provide broad spectrum resistance to viruses in insects: A comparative analysis of Wolbachia strains. PLOS Pathogens. 2014;10(9):e1004369. doi: 10.1371/journal.ppat.1004369.

26. Tain LS, Sehlke R, Meilenbrock RL, Leech T, Paulitz J, Chokkalingam M, et al. Tissue-specific modulation of gene expression in response to lowered insulin signalling in Drosophila. eLife. 2021;10:e67275. doi: 10.7554/eLife.67275.

27. Li J, Mahajan A, Tsai MD. Ankyrin repeat: a unique motif mediating protein-protein interactions. Biochemistry. 2006;45(51):15168–78. doi: 10.1021/bi062188q. PubMed PMID: 17176038.

28. Al-Khodor S, Price CT, Kalia A, Abu Kwaik Y. Functional diversity of ankyrin repeats in microbial proteins. Trends in Microbiology. 2010;18(3):132–9. doi: 10.1016/j.tim.2009.11.004.

29. Eichinger V, Nussbaumer T, Platzer A, Jehl MA, Arnold R, Rattei T. EffectiveDB--updates and novel features for a better annotation of bacterial secreted proteins and Type III, IV, VI secretion systems. Nucleic Acids Res. 2016;44(D1):D669-74. Epub 20151120. doi: 10.1093/nar/gkv1269. PubMed PMID: 26590402; PubMed Central PMCID: PMCPMC4702896.

30. Lamiable O, Kellenberger C, Kemp C, Troxler L, Pelte N, Boutros M, et al. Cytokine Diedel and a viral homologue suppress the IMD pathway in Drosophila. Proceedings of the National Academy of Sciences. 2016;113(3):698–703. doi: doi:10.1073/pnas.1516122113.

31. Goto A, Okado K, Martins N, Cai H, Barbier V, Lamiable O, et al. The kinase IKKβ regulates a STING- and NF-κB-dependent antiviral response pathway in Drosophila. Immunity. 2018;49(2):225–34.e4. doi: 10.1016/j.immuni.2018.07.013.

32. Liu Y, Gordesky-Gold B, Leney-Greene M, Weinbren NL, Tudor M, Cherry S. Inflammation-induced, STING-dependent autophagy restricts Zika virus infection in the Drosophila brain. Cell Host & Microbe. 2018;24(1):57–68.e3. Epub 2018/06/24. doi: 10.1016/j.chom.2018.05.022. PubMed PMID: 29934091; PubMed Central PMCID: PMCPMC6173519.

33. Palmer WH, Medd NC, Beard PM, Obbard DJ. Isolation of a natural DNA virus of Drosophila melanogaster, and characterisation of host resistance and immune responses. PLOS Pathogens. 2018;14(6):e1007050. doi: 10.1371/journal.ppat.1007050.

34. Dushay MS, Asling B, Hultmark D. Origins of immunity: Relish, a compound Rel-like gene in the antibacterial defense of Drosophila. Proceedings of the National Academy of Sciences. 1996;93(19):10343–7. doi: doi:10.1073/pnas.93.19.10343.

35. Meng X, Khanuja BS, Ip YT. Toll receptor-mediated Drosophila immune response requires Dif, an NF-kappaB factor. Genes and Development. 1999;13(7):792–7. Epub 1999/04/10. doi: 10.1101/gad.13.7.792. PubMed PMID: 10197979; PubMed Central PMCID: PMCPMC316597.

36. Zaidman-Rémy A, Poidevin M, Hervé M, Welchman DP, Paredes JC, Fahlander C, et al. Drosophila immunity: analysis of PGRP-SB1 expression, enzymatic activity and function. PLOS One. 2011;6(2):e17231. Epub 2011/03/03. doi: 10.1371/journal.pone.0017231. PubMed PMID: 21364998; PubMed Central PMCID: PMCPMC3041801.

37. Cherry S, Doukas T, Armknecht S, Whelan S, Wang H, Sarnow P, et al. Genome-wide RNAi screen reveals a specific sensitivity of IRES-containing RNA viruses to host translation inhibition. Genes Dev. 2005;19(4):445–52. doi: 10.1101/gad.1267905. PubMed PMID: 15713840; PubMed Central PMCID: PMCPMC548945.

38. Mizielinska S, Gronke S, Niccoli T, Ridler CE, Clayton EL, Devoy A, et al. C9orf72 repeat expansions cause neurodegeneration in Drosophila through arginine-rich proteins. Science. 2014;345(6201):1192–4. Epub 20140807. doi: 10.1126/science.1256800. PubMed PMID: 25103406; PubMed Central PMCID: PMCPMC4944841.

39. Paradkar PN, Trinidad L, Voysey R, Duchemin J-B, Walker PJ. Secreted Vago restricts West Nile virus infection in Culex mosquito cells by activating the Jak-STAT pathway. Proceedings of the National Academy of Sciences. 2012;109(46):18915–20.

40. Hedengren M, Dushay MS, Ando I, Ekengren S, Wihlborg M, Hultmark D. Relish, a central factor in the control of humoral but not cellular immunity in Drosophila. Molecular cell. 1999;4(5):827–37.

41. Kumar A, Srivastava P, Sirisena P, Dubey SK, Kumar R, Shrinet J, et al. Mosquito Innate Immunity. Insects. 2018;9(3). Epub 20180808. doi: 10.3390/insects9030095. PubMed PMID: 30096752; PubMed Central PMCID: PMCPMC6165528.

42. Lemaitre B, Hoffmann J. The host defense of Drosophila melanogaster. Annual Review of Immunology. 2007;25(2007):697–743. doi: 10.1146/annurev.immunol.25.022106.141615. PubMed PMID: WOS:000246437100024.

43. Myllymäki H, Valanne S, Rämet M. The Drosophila Imd signaling pathway. The Journal of Immunology. 2014;192(8):3455–62. doi: 10.4049/jimmunol.1303309.

44. Costa A, Jan E, Sarnow P, Schneider D. The Imd pathway is involved in antiviral immune responses in Drosophila. PLOS ONE. 2009;4(10):e7436. doi: 10.1371/journal.pone.0007436.

45. Kemp C, Mueller S, Goto A, Barbier V, Paro S, Bonnay F, et al. Broad RNA interference-mediated antiviral immunity and virus-specific inducible responses in Drosophila. Journal of Immunology. 2013;190(2):650–8. Epub 2012/12/21. doi: 10.4049/jimmunol.1102486. PubMed PMID: 23255357; PubMed Central PMCID: PMCPMC3538939.

46. Gendrin M, Zaidman-Rémy A, Broderick NA, Paredes J, Poidevin M, Roussel A, et al. Functional analysis of PGRP-LA in Drosophila immunity. PLOS ONE. 2013;8(7):e69742. doi: 10.1371/journal.pone.0069742.

47. Deddouche S, Matt N, Budd A, Mueller S, Kemp C, Galiana-Arnoux D, et al. The DExD/H-box helicase Dicer-2 mediates the induction of antiviral activity in drosophila. Nat Immunol. 2008;9(12):1425–32. Epub 20081026. doi: 10.1038/ni.1664. PubMed PMID: 18953338.

48. Mussabekova A, Daeffler L, Imler JL. Innate and intrinsic antiviral immunity in Drosophila. Cell and Molecular Life Sciences. 2017;74(11):2039–54. Epub 2017/01/20. doi: 10.1007/s00018-017-2453-9. PubMed PMID: 28102430; PubMed Central PMCID: PMCPMC5419870.

49. Dostert C, Jouanguy E, Irving P, Troxler L, Galiana-Arnoux D, Hetru C, et al. The Jak-STAT signaling pathway is required but not sufficient for the antiviral response of Drosophila. Nature Immunology. 2005;6(9):946–53. Epub 2005/08/09. doi: 10.1038/ni1237. PubMed PMID: 16086017.

50. Yamada R, Iturbe-Ormaetxe I, Brownlie JC, O’Neill SL. Functional test of the influence of Wolbachia genes on cytoplasmic incompatibility expression in Drosophila melanogaster. Insect Molecular Biology. 2011;20(1):75–85. Epub 2010/09/22. doi: 10.1111/j.1365-2583.2010.01042.x. PubMed PMID: 20854481.

51. Georgel P, Naitza S, Kappler C, Ferrandon D, Zachary D, Swimmer C, et al. Drosophila immune deficiency (IMD) is a death domain protein that activates antibacterial defense and can promote apoptosis. Developmental Cell. 2001;1(4):503–14. doi: 10.1016/S1534-5807(01)00059-4.

52. Adamo SA. Comparative Psychoneuroimmunology: Evidence From the Insects. Behavioral and Cognitive Neuroscience Reviews. 2006;5(3):128–40. doi: 10.1177/1534582306289580. PubMed PMID: 16891555.

53. Adelman JS, Martin LB. Vertebrate sickness behaviors: Adaptive and integrated neuroendocrine immune responses. Integrative and Comparative Biology. 2009;49(3):202–14. doi: 10.1093/icb/icp028.

54. Yamashita K, Oi A, Kosakamoto H, Yamauchi T, Kadoguchi H, Kuraishi T, et al. Activation of innate immunity during development induces unresolved dysbiotic inflammatory gut and shortens lifespan. Disease Models & Mechanisms. 2021;14(9). doi: 10.1242/dmm.049103.

55. Kurz CL, Charroux B, Chaduli D, Viallat-Lieutaud A, Royet J. Peptidoglycan sensing by octopaminergic neurons modulates Drosophila oviposition. eLife. 2017;6:e21937. doi: 10.7554/eLife.21937.

56. Dus M, Lai JS, Gunapala KM, Min S, Tayler TD, Hergarden AC, et al. Nutrient sensor in the brain directs the action of the bain-gut axis in Drosophila. Neuron. 2015;87(1):139–51. Epub 2015/06/16. doi: 10.1016/j.neuron.2015.05.032. PubMed PMID: 26074004; PubMed Central PMCID: PMCPMC4697866.

57. Yang Z, Huang R, Fu X, Wang G, Qi W, Mao D, et al. A post-ingestive amino acid sensor promotes food consumption in Drosophila. Cell Research. 2018;28(10):1013–25. doi: 10.1038/s41422-018-0084-9.

58. Lee K-M, Daubnerová I, Isaac RE, Zhang C, Choi S, Chung J, et al. A neuronal pathway that controls sperm ejection and storage in female Drosophila. Current Biology. 2015;25(6):790–7. doi: 10.1016/j.cub.2015.01.050.

59. De Gregorio E, Spellman PT, Rubin GM, Lemaitre B. Genome-wide analysis of the Drosophila immune response by using oligonucleotide microarrays. Proceedings of the National Academy of Sciences. 2001;98(22):12590–5. doi: doi:10.1073/pnas.221458698.

60. Li X, Rommelaere S, Kondo S, Lemaitre B. Renal purge of hemolymphatic lipids prevents the accumulation of ROS-induced inflammatory oxidized lipids and protects Drosophila from tissue damage. Immunity. 2020;52(2):374–87.e6. Epub 2020/02/23. doi: 10.1016/j.immuni.2020.01.008. PubMed PMID: 32075729.

61. Ohhara Y, Kobayashi S, Yamakawa-Kobayashi K, Yamanaka N. Adult-specific insulin-producing neurons in Drosophila melanogaster. Journal of Comparative Neurology. 2018;526(8):1351–67. doi: 10.1002/cne.24410.

62. Walsh D, Mathews MB, Mohr I. Tinkering with translation: protein synthesis in virus-infected cells. Cold Spring Harbor Perspectives in Biology. 2013;5(1):a012351. Epub 2012/12/05. doi: 10.1101/cshperspect.a012351. PubMed PMID: 23209131; PubMed Central PMCID: PMCPMC3579402.

63. Grobler Y, Yun CY, Kahler DJ, Bergman CM, Lee H, Oliver B, et al. Whole genome screen reveals a novel relationship between Wolbachia levels and Drosophila host translation. PLOS Pathogens. 2018;14(11):e1007445. doi: 10.1371/journal.ppat.1007445.

64. Nagarajan S, Grewal SS. An investigation of nutrient-dependent mRNA translation in Drosophila larvae. Biology Open. 2014;3(11):1020–31. doi: 10.1242/bio.20149407.

65. Rances E, Ye YH, Woolfit M, McGraw EA, O’Neill SL. The relative importance of innate immune priming in Wolbachia-mediated dengue interference. PLoS Pathog. 2012;8(2):e1002548. Epub 20120223. doi: 10.1371/journal.ppat.1002548. PubMed PMID: 22383881; PubMed Central PMCID: PMCPMC3285598.

66. Rances E, Johnson TK, Popovici J, Iturbe-Ormaetxe I, Zakir T, Warr CG, et al. The toll and Imd pathways are not required for wolbachia-mediated dengue virus interference. J Virol. 2013;87(21):11945–9. Epub 20130828. doi: 10.1128/JVI.01522-13. PubMed PMID: 23986574; PubMed Central PMCID: PMCPMC3807350.

67. Kambris Z, Cook PE, Phuc HK, Sinkins SP. Immune activation by life-shortening Wolbachia and reduced filarial competence in mosquitoes. Science. 2009;326(5949):134–6. doi: 10.1126/science.1177531. PubMed PMID: 19797660; PubMed Central PMCID: PMCPMC2867033.

68. Moreira LA, Iturbe-Ormaetxe I, Jeffery JA, Lu G, Pyke AT, Hedges LM, et al. A Wolbachia symbiont in Aedes aegypti limits infection with dengue, Chikungunya, and Plasmodium. Cell. 2009;139(7):1268–78. doi: 10.1016/j.cell.2009.11.042. PubMed PMID: 20064373.

69. Meng-Yan Chen DL, Zhi-Ning Wang, Feng-Zhen Xu, Yi-Wei Feng, Qiong-Lin Yu, Ying-Ying Wang, Shu Zhang, Yu-Feng Wang. Infection by virulent wMelPop Wolbachia improves learning and memory capacity in Drosophila melanogaster,. Animal Behaviour. 2024;212:101–12.

70. Serbus LR, White PM, Silva JP, Rabe A, Teixeira L, Albertson R, et al. The impact of host diet on Wolbachia titer in Drosophila. PLoS Pathog. 2015;11(3):e1004777. Epub 20150331. doi: 10.1371/journal.ppat.1004777. PubMed PMID: 25826386; PubMed Central PMCID: PMCPMC4380406.

71. Ahlers LRH, Trammell CE, Carrell GF, Mackinnon S, Torrevillas BK, Chow CY, et al. Insulin potentiates JAK/STAT signaling to broadly inhibit Flavivirus replication in insect vectors. Cell Reports. 2019;29(7):1946–60.e5. doi: 10.1016/j.celrep.2019.10.029.

72. Wu M, Sun LV, Vamathevan J, Riegler M, Deboy R, Brownlie JC, et al. Phylogenomics of the reproductive parasite Wolbachia pipientis wMel: a streamlined genome overrun by mobile genetic elements. PLOS Biology. 2004;2(3):E69. Epub 2004/03/17. doi: 10.1371/journal.pbio.0020069. PubMed PMID: 15024419; PubMed Central PMCID: PMCPMC368164.

73. Rice DW, Sheehan KB, Newton ILG. Large-scale identification of Wolbachia pipientis effectors. Genome Biology and Evolution. 2017;9(7):1925–37. doi: 10.1093/gbe/evx139.

74. Pinto SB, Riback TIS, Sylvestre G, Costa G, Peixoto J, Dias FBS, et al. Effectiveness of Wolbachia-infected mosquito deployments in reducing the incidence of dengue and other Aedes-borne diseases in Niteroi, Brazil: A quasi-experimental study. PLoS Negl Trop Dis. 2021;15(7):e0009556. Epub 20210712. doi: 10.1371/journal.pntd.0009556. PubMed PMID: 34252106; PubMed Central PMCID: PMCPMC8297942.

75. Osterwalder T, Yoon KS, White BH, Keshishian H. A conditional tissue-specific transgene expression system using inducible GAL4. Proceedings of the National Academy of Sciences. 2001;98(22):12596–601. Epub 2001/10/25. doi: 10.1073/pnas.221303298. PubMed PMID: 11675495; PubMed Central PMCID: PMCPMC60099.

76. Bass TM, Weinkove D, Houthoofd K, Gems D, Partridge L. Effects of resveratrol on lifespan in Drosophila melanogaster and Caenorhabditis elegans. Mechanisms of Ageing and Development. 2007;128(10):546–52. doi: 10.1016/j.mad.2007.07.007.

77. Bischof J, Maeda RK, Hediger M, Karch F, Basler K. An optimized transgenesis system for Drosophila using germ-line-specific φC31 integrases. Proceedings of the National Academy of Sciences. 2007;104(9):3312–7. doi: doi:10.1073/pnas.0611511104.

78. Grabherr MG, Haas BJ, Yassour M, Levin JZ, Thompson DA, Amit I, et al. Full-length transcriptome assembly from RNA-Seq data without a reference genome. Nat Biotechnol. 2011;29(7):644–52. Epub 20110515. doi: 10.1038/nbt.1883. PubMed PMID: 21572440; PubMed Central PMCID: PMCPMC3571712.

79. Li B, Dewey CN. RSEM: accurate transcript quantification from RNA-Seq data with or without a reference genome. BMC Bioinformatics. 2011;12:323. Epub 20110804. doi: 10.1186/1471-2105-12-323. PubMed PMID: 21816040; PubMed Central PMCID: PMCPMC3163565.

80. Wong R, Piper MDW, Wertheim B, Partridge L. Quantification of food intake in Drosophila. PLOS ONE. 2009;4(6):e6063. doi: 10.1371/journal.pone.0006063.

81. Ritchie ME, Phipson B, Wu D, Hu Y, Law CW, Shi W, et al. limma powers differential expression analyses for RNA-sequencing and microarray studies. Nucleic Acids Research. 2015;43(7):e47. Epub 2015/01/22. doi: 10.1093/nar/gkv007. PubMed PMID: 25605792; PubMed Central PMCID: PMCPMC4402510.

82. Ge SX, Jung D, Yao R. ShinyGO: a graphical gene-set enrichment tool for animals and plants. Bioinformatics. 2019;36(8):2628–9. doi: 10.1093/bioinformatics/btz931.

83. Longdon B, Hadfield JD, Day JP, Smith SCL, McGonigle JE, Cogni R, et al. The causes and consequences of changes in virulence following pathogen host shifts. PLOS Pathogens. 2015;11(3):e1004728. doi: 10.1371/journal.ppat.1004728.

84. Zhang P, Catterson JH, Gronke S, Partridge L. Inhibition of S6K lowers age-related inflammation and increases lifespan through the endolysosomal system. Nat Aging. 2024. Epub 20240227. doi: 10.1038/s43587-024-00578-3. PubMed PMID: 38413780.

